# Spatio-temporal coordination of the DNA Double-Strand Break repair machinery in *Escherichia coli*

**DOI:** 10.1101/2024.12.18.629153

**Authors:** Daniel Thédié, Alessia Lepore, Lorna McLaren, Meriem El Karoui

## Abstract

Repairing DNA damage is of primary importance for all living organisms. DNA double-strand breaks (DSBs) are one of the most serious types of DNA damage, as they lead to loss of genetic information and death when not repaired. In *E. coli*, they are recognised and processed by the RecBCD complex, which initiates repair by homologous recombination. Although the repair dynamics down-stream of RecBCD has been well characterised, it is still unclear how long this complex stays attached to DNA and what triggers its dissociation *in vivo*. To answer these questions, we imaged RecB at the single-molecule level, and quantified its dynamic behaviour in bacterial cells exposed to ciprofloxacin, an antibiotic that induces DSBs. Our results show that RecB forms long-lived complexes with DSBs (10 seconds), and that its dissociation from DNA is an intrinsic property of the complex, that does not depend on the amount of DNA damage, nor the following steps in the repair pathway. Moreover, we show that we can use RecB binding to DSBs as a marker to estimate the rate of damage formation. This study provides a detailed quantitative insight into the interaction of RecBCD with DNA double-strand ends in *E. coli in vivo*, and into the bacterial response to DSBs induced by ciprofloxacin.

## Introduction

Repairing DNA damage and ensuring the integrity of the genome is essential to all living organisms. Among the different types of damage, DNA double-strand breaks (DSBs) pose a serious threat to bacterial cells: when left unrepaired, they can lead to cell death, genome rearrangements or mutagenesis (1). In *E. coli*, DSBs are repaired through the RecBCD pathway. RecBCD is a heterotrimeric protein complex, which combines several enzymatic activities (2). Upon recognition of a DSB, RecBCD translocates along the DNA at a speed of ∼1.6 kb/s (3), and digests both DNA strands until it recognises a specific 8-base pair sequence (5’-GCTGGTGG-3’) called a Chi-site. RecBCD then stops degrading the 3’ end, leading to the formation of a 3’-OH DNA overhang. Chi recognition by RecBCD has been previously reported to occur with ∼40% probability (4, 5). As RecBCD creates the 3’ overhang, the RecB subunit promotes loading of the RecA protein onto single-stranded DNA (ssDNA) (6, 7). The resulting RecA-DNA nucleoprotein filament is used as a template to search for an intact homologous sequence and perform homologous recombination (8, 9). It has been suggested that RecBCD dissociates from DNA by disassembly of the three subunits following Chi recognition (10); however, it is still unclear what specifically triggers the dissociation and how long RecBCD stays bound to DNA before dissociating.

A common source of DSBs in *E. coli* is the collision of the replication fork with DNA-bound proteins, which leads to the disassembly of the replisome (11). It has been reported that ∼18% of cells undergo endogenous DNA damage per cell cycle, most likely from replication fork breakage (12). Replisome disassembly results in a Y-shaped DNA structure that is bound by the RuvAB complex, which pulls the DNA strands (in a process called “branch migration”) to create a “chicken-foot” four-way DNA structure (13). The free end of this structure is a DNA double strand end, and is bound by RecBCD, which either (i) digests the whole DNA tail and displaces RuvAB, allowing the replisome to be re-loaded in a PriA-dependent manner (13), or (ii) recognises a Chi-site and loads the RecA protein onto the 3’ end of the DNA tail, leading to SOS induction (14). Since this part of the chromosome was recently replicated, a homologous sequence is likely to be present in close proximity, and can be used for homologous recombination.

*E. coli* cells might also experience DSBs from exogenous sources, such as antibiotics. Ciprofloxacin is an antibiotic of the fluoroquinolones family, which causes DSBs. In *E. coli*, it does so by binding covalently to the topoisomerase II DNA gyrase (and with lower affinity to topoisomerase IV), and trapping it in a DNA-bound conformation (15). The exact mechanism through which DSBs are created is still partially unclear, but ciprofloxacin has been suggested to cause replication-dependent and independent DSBs (16). Since poisoned gyrase is closely bound to DNA, it is likely to cause replication fork collisions (17, 18), and the subsequent formation of a chicken foot structure. Furthermore, the ciprofloxacin-poisoned gyrase is trapped in the “open” stage of its catalytic cycle, where it holds together two disjointed DNA ends. Therefore, after gyrase removal through a process that involves ExoVII nuclease (19), it is expected that a DSB might be formed independently of DNA replication (20). DSB repair is the first step of a general response to DNA damage known as the SOS response (21). The formation of the RecA nucleoprotein filament triggers the auto-proteolysis of the LexA transcriptional repressor, which results in the activation of the genes of the SOS regulon. Among those are RecA itself, inhibitors of cell division such as SulA, and the SMC (Structural Maintenance of Chromosomes)-like protein RecN. As a result, the cells elongate (22), and their DNA is compacted at the cell centre (23).

Mutations in proteins of the repair pathway have varying effects on the cell’s ability to repair DSBs. Cells that are *recA*-deficient (Δ*recA*) are entirely unable to repair DSBs through homologous recombination, and the prolonged action of RecBCD on the DNA results in the digestion of large parts of the chromosome (24, 25). *recA*-deficient cells are, however, capable of processing DSBs arising from fork reversal, thanks to RecBCD’s exonuclease activity (13, 14). This enables *recA*-deficient cells to repair replication-induced DNA damage, resulting in a growth rate in the absence of DNA damage close to that of wild-type cells (26). The *D*_1080_ → *A* point mutation in the RecB subunit of the RecBCD complex (RecB_1080_) strongly reduces RecB’s nuclease activity, as well as its ability to load the RecA protein on DNA (27, 28). The other activities of the complex (DSB recognition, DNA unwinding, Chi-recognition) are unaffected (29). Despite the loss of its RecA loading activity, the *recB*_1080_ mutant is still able to repair DSBs. To do so, the RecB_1080_CD complex unwinds DNA at the break without degrading it, and the RecJ exonuclease degrades the 5’ ssDNA. Finally, the RecFOR complex loads RecA, allowing for homologous recombination to occur (30).

Previous microscopy studies have shed light on different aspects of DSB repair by RecBCD. Several *in vitro* studies have quantified the rate of DNA unwinding by RecBCD (31, 32), as well as the growth rate of RecA filaments (33–35) and the mechanism of homology search (36, 37). Complementary to the *in vitro* work, *in vivo* studies have provided insight into DNA end resection by RecBCD (3), confirming its high translocation speed and processivity. More recently, we investigated RecB mobility *in vivo* at the single-molecule level (38), finding that RecB’s engagement in repair initiation is proportional to the level of DNA damage, and that its interactions with DNA are consistent with its role in initiating the repair process. Furthermore, we observed that the efficiency of DNA repair depends on the full functionality of RecB’s activities. The spatial organisation of RecA during DSB repair has been extensively characterised. Studies reported the presence of RecA foci (39– 42), filaments (43), or bundles of filaments (44, 45). The homology search and pairing processes were also observed *in vivo*, showing that chromosomes are brought to the centre of the cell during the repair process (9, 46). Our previous observations (38) highlighted how the coordination of all RecBCD activities facilitates rapid and efficient repair. Nevertheless, several important aspects of RecBCD’s processing of DSBs *in vivo* remain to be clarified: how long does RecBCD stay bound to DNA after recognising a DSB? Do the recruitment and binding pattern change under different levels of DNA damage? How does its spatio-temporal organisation change upon initiation of DNA damage? Does RecA play a role in triggering RecBCD dissociation from DNA? These open questions are fundamental to the complete understanding of DSB repair in *E. coli*.

To address these questions, we used single-molecule imaging to quantify RecB binding to DSBs in living *E. coli* cells exposed to ciprofloxacin. We measured the duration of the RecB-DSB interaction and found that it was not affected by the concentration of ciprofloxacin. Concurrent imaging of RecB and the bacterial nucleoid showed compaction of the bacterial nucleoid, which was associated with RecB’s subsequent recruitment to new DSBs. Additional imaging of RecA revealed structures similar to previously observed bundles, which scaled with ciprofloxacin concentration and duration of exposure. Furthermore, we imaged RecB in the Δ*recA* and *recB*_1080_ mutants, and revealed that RecB dissociation depends only on the intrinsic biochemical activities of the complex. These results reveal the importance of the intricate spatio-temporal coordination of the DSB repair machinery to ensure efficient repair in *E. coli*.

## Materials and Methods

### Strain construction

*E. coli* MG1655 and derivatives were used in this study. A list of all strains and plasmids used are presented in Supp. Tables 1 and 2. MEK2623 (RecB-Halo RecA-SYFP2) was constructed by transferring a RecA-SYFP2 tandem fusion by P1 transduction, using EL2514 (9) as a donor and MEK65 (47) as a receiver. The clones were selected on kanamycin plates and checked by Sanger sequencing. MEK2622 (Δ*recA*) was constructed by transferring the *recA* deletion by P1 transduction, using DL654 as a donor, and MEK65 as a receiver. MEK2629 was obtained by transforming the pDT6 plasmid (a gift from Prof. Mark Dillingham) into the MEK65 strain by heat shock.

### Microscopy samples preparation

For all experiments, the cells were grown in M9 supplemented with 0.2% (w/v) glucose, 2 mM MgSO_4_, 0.1 mM CaCl_2_, 1X MEM Essential and MEM non-essential amino acids (Gibco).

#### Cell culture

Cells were grown overnight from a −80°C glycerol stock in 5 mL medium at 37°C, shaking at 150 rpm. In the morning, the cells were diluted 1:300 into 15 mL of medium and grown at 37°C, shaking at 150 rpm until they reached an OD_600_ of 0.2–0.3 (mid-exponential phase). Alternatively, a 15 mL culture was inoculated using a −80°C glycerol stock, and placed at 37°C with stirring in a turbidostat device (OGIBio), which measured OD_600_ and diluted cells at 3 min intervals, to keep them at a constant OD_600_ of 0.2.

#### Halo labelling

We used a labelling protocol similar to our previous study (38). A volume of cells equivalent to 1 mL at OD_600_=0.2 was centrifuged (8000 rpm) and resuspended in 1 mL of fresh medium. 5 µL of JF549 dye (Janelia Fluor Halo-tag Ligand, Promega, resuspended in DMSO) was added (final concentration 1 µM) and the cells were further incubated for 1h at 37°C in the dark, shaking at 150 rpm.

#### DNA labelling

When relevant, cellular DNA was labelled by adding Sytox Green (Invitrogen, catalog number S7020) at a concentration of 500 nM, at the same time as the JF549 dye using similar incubation conditions (1h at 37°C in the dark, shaking at 150 rpm).

#### Dye removal

The same procedure was followed to remove unbound JF549 and Sytox Green dye. The cells were centrifuged for 3 min at 8000 rpm, the supernatant discarded, and the pellet resuspended in 1 mL of fresh medium and transferred to a new tube to avoid the dyes sticking to the plastic of the tube. This procedure was repeated 3 times to fully remove unbound dye. The cells were finally resuspended in 200 µL medium.

#### Gam overexpression

For experiments where Gam was over-expressed, arabinose was added to the cells during the labelling step, at 1% (w/v) final concentration. Arabinose (at the same concentration) was also added to the medium for subsequent washes, and in the agarose pad to maintain exposure during imaging.

#### Sample preparation

Agarose pads were prepared by dissolving 2% (w/v) agarose in culture medium. In experiments with ciprofloxacin, the antibiotic was added to the agarose pad at the desired final concentration. After the dye removal step, 5 µL of cells were added on the agarose pad, and left to settle for 10 min at 37°C in the dark before imaging.

### Microscopy

#### Microscope setup

Imaging was performed using an inverted microscope (Nikon Ti-E) equipped with a 100X TIRF Nikon objective (NA 1.49, oil immersion) and a 1.5X Nikon magnification lens (pixel size = 107 nm). Fluorescence excitation was achieved using 488- and 561-nm lasers (Coherent OBIS) in Highly Inclined Laminar Optical sheet (HILO) configuration. Excitation light and fluorescence emission were separated using a dual-wavelength dichroic filter (TRF59904, Chroma), and the fluorescence signal was detected on an Electron Multiplying Charge-Coupled Device (EMCCD) camera (iXion Ultra 897, Andor). The hardware was controlled and images were saved using MetaMorph (Molecular Devices; v7.8.13.0). The HILO configuration was established using the iLas variable angle TIRF control window (Gattaca), with an angle of 57°. All experiments were performed at 37°C, using an Okolab microscope cage incubator equipped with dark panels.

#### Data acquisition

For each sample, a Metamorph journal was used to acquire images on 40 different positions, each separated by 150 µm. The full camera sensor (512 × 512 pixels) was used. Acquisition parameters are described in Supp. Table 3. Brightfield images were acquired in all experiments, as well as two fluorescence channels: JF549 (RecB) and either SYFP2 (RecA) or Sytox Green (nucleoid). At each position, 50 images were acquired at 2-second intervals for RecB and RecA, or a single image was acquired for the nucleoid (Supp. Fig. 1). The prolonged imaging of the RecB-Halo fusion was only possible thanks to the exceptional photostability of the JF549 dye, which displayed slow photobleaching and no blinking (Supp. Note 1 and Supp. Fig. 2).

### Data analysis

The general data analysis workflow is described in Supp. Figure 3. In brief, after acquisition, the raw microscopy images were stored in `.tif` format on a dedicated Omero server (OME). They were directly accessed by the BACM-MAN ImageJ plugin (48), which performed image processing tasks such as denoising of fluorescence images, cell segmentation (including manual curation of incorrectly segmented cells), nucleoid segmentation, and fluorescent spot detection. Measurement tables in CSV format were exported from BACMMAN and loaded in Jupyter Notebooks using the PyBerries package (version 0.2.25) in Python (version 3.11). The data tables were manipulated using PyBerries and the pandas library (version 2.0.2). Figures were generated in Jupyter Notebooks using the Seaborn library (version 0.13.2). Figures that contained several panels were assembled in Inkscape (version 1.3.2).

### BACMMAN pipeline

#### Deep-learning denoising of RecB-Halo fluorescence images

We improved a self-supervised denoising neural network by making it use the previous and next frames in the timelapse and performing deconvolution simultaneously. Our method is based on a previously developed method (49) which uses an encoder-decoder convolutional neural network that performs denoising (DNet), as well as a fully connected neural network that recovers the noise distribution used to compute the self-supervised loss. In order to include previous and next frames, we modified the architecture of DNet so that the encoder is shared between frames (t-1, t and t+1), i.e. it processes each frame independently. At each contraction level, the residual tensors corresponding to each frame are combined using a 1× 1 convolution and fed to the decoder. The same applies for the feature tensors and the last level of the decoder. The decoder only predicts the central frame (t) and the same self-supervised loss is used. We observe a dramatic improvement of denoising performance, which implies that the proposed architecture benefits from using information from adjacent frames. It has been shown that performing denoising and deconvolution simultaneously dramatically improves deconvolution performance (50). We computed the experimental PSF of our system by averaging hundreds of bead images cropped in a 33 × 33 pixels area around each bead, and normalised so that values sum to one. The obtained averaged images were used during training as kernel for a 2D convolution at the end of the neural network. At prediction time, convolution was removed to obtain a deconvolved image.

#### Cell segmentation

The 16-image brightfield Z-stack was first cropped to 5 images on one side of the focus, as required by our segmentation algorithm. The 5-image stack was used as input to Talissman, a U-net-based segmentation algorithm. In brief, the U-net model predicts a Euclidean distance map, where the value of each pixel is its predicted distance to the nearest background pixel. A watershed algorithm is then applied to retrieve cell contours. This approach allowed us to accurately segment cells from bright-field images, including when they formed tight clusters. Any overlapping cells were manually removed from the analysis. Following segmentation, post-filters were applied to dilate the segmented regions slightly and to remove any cells that were in contact with the edge of the image and might, therefore, be cropped. Finally, all segmentation masks were visually inspected and curated to remove cells that were incorrectly segmented (∼1% of total cells).

#### Detection of RecB-Halo spots

Fluorescent spots were segmented using a seeded watershed algorithm on the Laplacian transform of the denoised RecB image. The quality of the segmentation was visually assessed by overlaying the segmentation mask on the raw fluorescence images. The segmentation parameters were optimised by trial and error to minimise the error rate (assessed by visual inspection of raw fluorescence images overlaid with the segmentation masks), but no manual curation was applied to avoid introducing user bias in the results.

#### Classification of RecA structures

We designed a U-net-based deep-learning model capable of classifying objects (in this case, cells) based on the type of RecA structure they contained (Supp.Figure 4). The model was trained on a dataset of 4,069 cells, manually labelled as containing either a RecA focus, a RecA filament, or neither of those (homogenous diffuse fluorescence). Because there were different numbers of each predicted class in the training set, the loss function (categorical cross-entropy) was weighted by the inverse class frequency (weights were 21, 101 and 133 for the diffuse, filament and focus classes, respectively). The model was trained on 128 × 128 pixel crops of the RecA-SYFP2 fluorescence images, with the corresponding cell segmentation mask as input. Data augmentation was performed using the [dataset iterator](https://github.com/jeanollion/dataset_iterator) package by applying random rotations, flips, and elastic deformations. The model was trained for 1000 epochs with a batch size of 3, using the Adam optimiser. The learning rate was initially set to 0.0004, and reduced by a factor or 10 when the validation loss did not improve for 30 consecutive epochs (to a minimum of 1 ×10^−6^). The model’s predictions were evaluated against a test set of 15 manually-labelled images that were not used in training, achieving an accuracy of 84%.

#### Nucleoid segmentation

Sytox Green fluorescence images were processed with a rolling-ball noise subtraction (radius 6 pixels), and nucleoids segmented by a seeded watershed algorithm. The seed threshold was determined by Otsu’s method (51). Any segmented regions that were in contact with each other within the same cell were merged, and segmented regions smaller than 5 pixels were excluded. Regions with weak signal-to-noise ratio (SNR < 2) were also excluded. For each segmented region, the SNR was computed as follows:

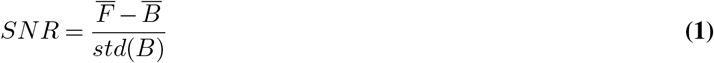

with 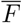 the mean fluorescence intensity in the object, 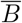 the mean background intensity, and *std*(*B*) the standard deviation of background intensities.

#### Measurements

As the final step of the pipeline, BACMMAN performed several measurements on the objects created (cells, RecB spots and nucleoids). These measurements included cell length, cell area, raw fluorescence intensity, background-subtracted fluorescence intensity, number of RecB spots in the cell, and position of the RecB spots and nucleoids along the long and short axis of the cell. For each dataset, BACMMAN produced one csv table per object.

### PyBerries

#### Data import and format

The PyBerries package, combined with the pandas library, enabled easy import of the multiple CSV tables produced by BACMMAN, and performing operations such as grouping, aggregations, normalisation and curve fitting.

#### Fitting Halo-RecB spot lifetimes

The lifetime of individual RecB spots was computed in BACMMAN. The resulting lifetime histogram was fitted with a bi-exponential decay function of the 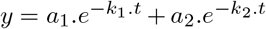. To improve the reliability of the fits, a bootstrapping procedure was used. For each ciprofloxacin concentration, an array of RecB spot lifetimes of the same size as the original data was drawn with replacement (i.e. each lifetime could potentially be drawn several times), and the resulting histogram was fitted with the bi-exponential decay model. This procedure was repeated 100 times, yielding 100 different estimates for each fit parameter. The best fit parameter was obtained by taking the median of all estimates.

Because it is not possible to differentiate whether the disappearance of a fluorescent spot was a result of photobleaching or of unbinding and returning to the pool of diffusing molecules, we considered that the fitted “spot disappearance rates” *k*_1_ and *k*_2_ were a sum of RecB’s dissociation rate from DNA and the dye’s bleaching rate (*k*_1_ = *k*_*d*1_ + *k*_*b*_ and *k*_2_ = *k*_*d*2_ + *k*_*b*_, with *k*_*d*1_ and *k*_*d*2_ the two RecB dissociation rates, and *k*_*b*_ the bleaching rate). The bleaching rate was calculated for each dataset, as described in Supp. Note 1 and Supp. Figure 2, and subtracted to retrieve the true RecB dissociation rates from DNA.

#### Estimation of the rate of RecB recruitment to DSBs

For each timelapse, we quantified the total number of RecB spots that were bound to DSBs. This could not be achieved by thresholding of the lifetime histograms, as a significant number of short-lived spots may correspond to DSB-bound RecB molecules. To estimate the total number of DSB-bound RecB, we considered that all RecB spots that disappeared with a slower rate (as determined by the bi-exponential decay fit) were DSB-bound. We multiplied the total number of spots that appeared during the timelapse by the proportion of slow-dissociating spots retrieved from the lifetime histogram fits, which gave us an estimate of the number of DSB-bound RecB per timelapse, from which a number of recruitment events per cell per hour was calculated. This rate was corrected for photobleaching of the fluorescent dye, to account for non-fluorescent RecBCD complexes binding to DSBs. This was done independently for each dataset, by computing the average proportion of the remaining initial fluorescence over time. The number of RecB binding events recorded was divided by this average proportion of fluorescence to estimate the real number of RecB binding events. For example: for 2 RecB recruitments per cell per hour observed, and an average of 50% of the initial fluorescence remaining over the timelapse, the estimated number of RecB recruitments per cell per hour would be 2*/*0.5 = 4. This estimation method assumes that RecB binding events are equally likely at all points of the timelapse.

### Code availability

Instructions to install the BACMMAN ImageJ plugin can be found on the [BACMMAN wiki](https://github.com/jeanollion/bacmman/wiki/Installation). The deep-learning models used for cell segmentation, fluorescence denoising and RecA structure classification can be found in BACMMAN’s deep-learning models library. To access the library, in BACMMAN go to the Import/Export menu and select Online DL Model library. Under Username enter “el-karoui-lab”, leave the password field empty and click on Connect. The models can be found in the “Thedie-et-al” folder. BACMMAN dataset configurations can be found in the [BACMMAN configuration library](https://github.com/jeanollion/bacmman/wiki/Online-Configuration-Library). To access the library, in BACMMAN go to the Import/Export menu and select Online Configuration Library. Under Username enter “el-karoui-lab”, leave the password field empty and click on Connect. The configurations can be found in the “Thedie-et-al” folder. The PyBerries package can be found on the [Python Package Index](https://pypi.org/project/PyBerries/), and its source code is available on [Gitlab](https://gitlab.com/MEKlab/bacmman-object-classifier). The deep-learning model used to classify cells according to their RecA-associated fluorescence is available on Gitlab. The Jupyter Notebooks used to make the article figures are available on [Github](https://github.com/DanielThedie/RecB_article).

## Results

### RecB forms long-lived spots when recruited to DSBs caused by ciprofloxacin

To quantify RecB binding time to DSBs in live *E. coli*, we used a Halo-tag fusion to the RecB subunit, conjugated to the JF549 fluorescent dye (Figure 1A). The fusion was previously fully characterised, ensuring specific one-to-one labelling of RecB molecules without adverse effects on the DNA repair process (38, 47). As a result of the large size and topological constraints of the bacterial chromosome, DSB-bound RecB diffuses very slowly (38). Therefore, we applied the previously developed technique of localisation enhancement (52, 53): since RecB is present at low copy numbers in *E. coli* [∼5 molecules per cell on average (47, 54)], imaging live cells with a long exposure time (1 second) made fast-diffusing RecB molecules appear as weak homogeneous signal in the cell, while very slow-diffusing RecB molecules formed diffraction-limited spots (referred to throughout this article as RecB spots, Figures 1B and 1C). The complete absence of similar spots in cells expressing the free Halo-tag from a plasmid confirmed that these spots were specific to RecB (Supp. Figure 5). However, their exact nature needed to be confirmed. Although DSB-bound molecules are expected to diffuse very slowly, the RecBCD-Halo complex diffuses in the crowded and constrained environment of the cytoplasm, and might undergo transient interactions with DNA (38), which could lead to RecB spots forming independently of DSB binding. To gain more insight into the nature of RecB spots, we increased the number of DSBs by adding ciprofloxacin to agar pads before imaging, and measured the lifetime of the spots by recording 100-second timelapse videos with 2-sec interval between frames at 40 different positions in the sample (Figure 2A). The total duration of the experiment was 75 min following exposure to ciprofloxacin. The 1-second exposure time was optimised to detect RecB spots with good signal-to-noise while limiting dye photobleaching and keeping a sufficient temporal resolution to determine spot lifetimes. Visual inspection of the timelapse data showed that RecB spot lifetime was hetereogenous, with a majority of spots being visible only for 1–2 frames, and a small number persisting for longer times (Figure 2A). These longer-lived spots became more frequent as the ciprofloxacin concentration increased. Background-subtracted intensity time-traces for single RecB spots showed that spots lost their intensity in a single step (Supp. Figure 6), consistent with single-molecule photobleaching or transitioning from a very slowly diffusing state to a rapid one. Therefore, our imaging setup allowed us to visualise single, very slow diffusing RecB molecules, and to measure the time they spent in this diffusive state.

**Fig. 1.**
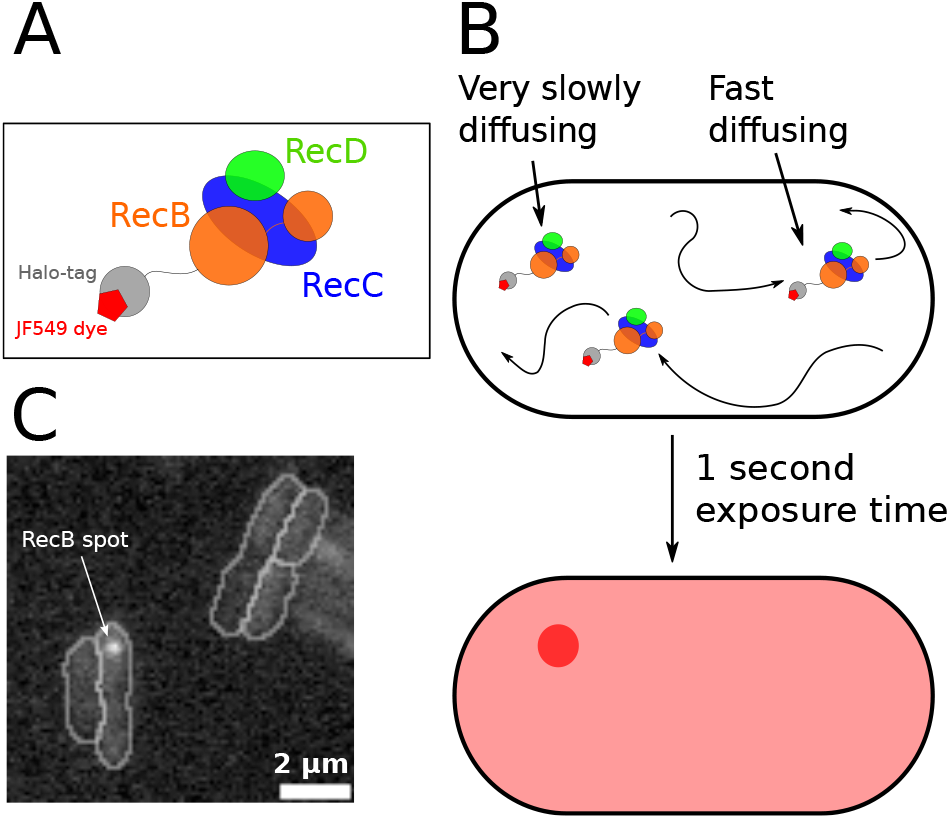
Detecting RecB binding to DSBs. **(A)** Scheme of our experimental protocol. The RecB subunit of the RecBCD complex is fused to a Halo-tag, bound by the JF549 fluorescent dye (38, 47). **(B)** A long exposure time (1 sec) makes diffusing molecules appear as a diffuse signal in the cell, while DSB-bound molecules are visible as bright, diffraction-limited spots. **(C)** Example image of a RecB spot (white arrow).

**Fig. 2.**
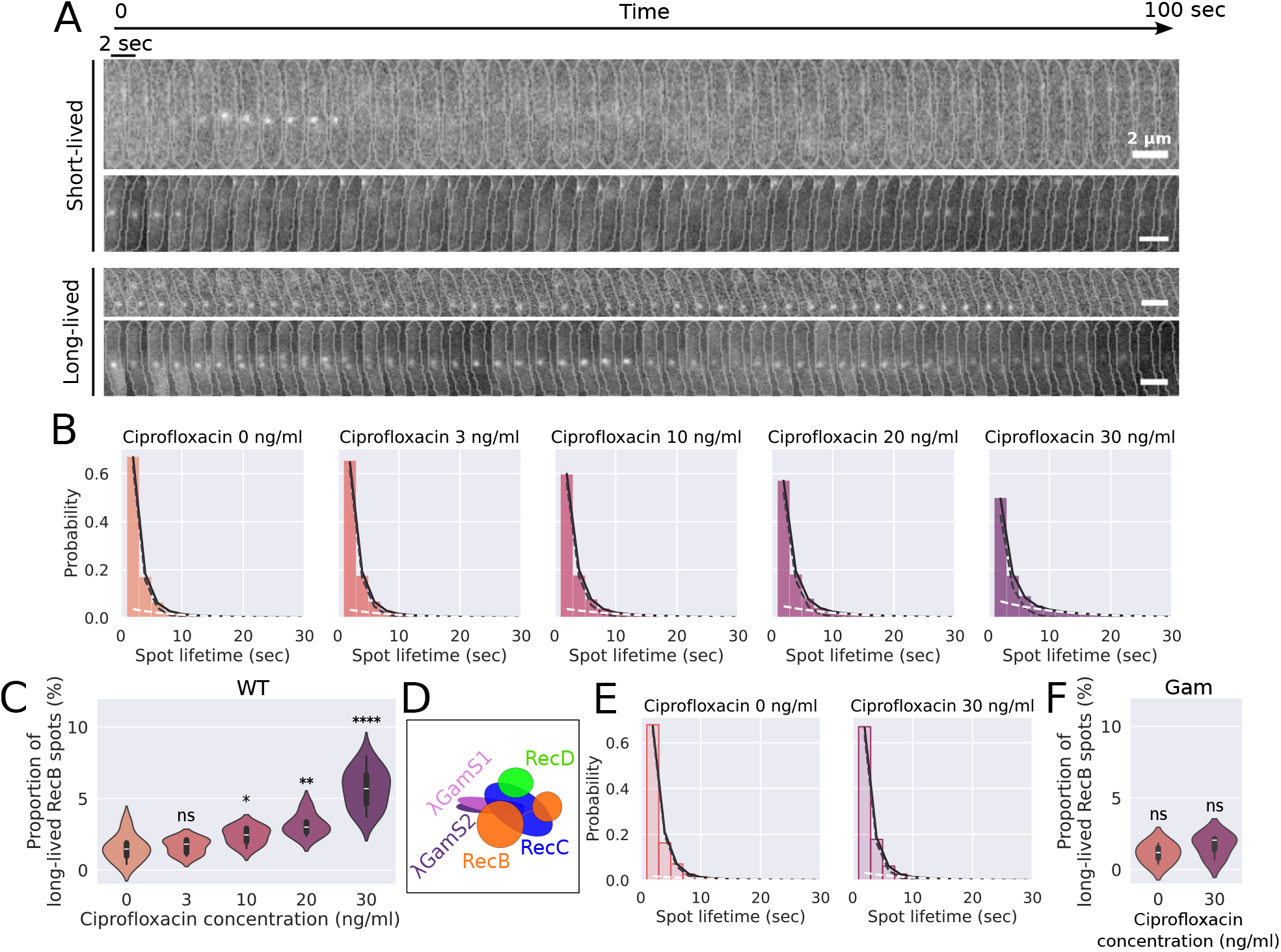
RecB spot lifetimes under ciprofloxacin exposure. **(A)** Example kymographs (2-sec interval, 100 sec total) of cells containing short- and long-lived RecB spots. **(B)** RecB spot lifetime histograms at 0, 3, 10, 20 and 30 ng/ml ciprofloxacin (bars), fitted with a bi-exponential decay model (black line, fit components shown as dashed lines). N_cells_ = 66,764. N_spots_ = 170,138. **(C)** Proportion of long-lived RecB spots as a function of ciprofloxacin concentration for wild-type (WT) cells. Violin plots show single-cell distributions, box plots show the median and interquartile range. Statistically significant differences to the WT, no ciprofloxacin condition (Mann-Whitney test) are shown as follows: ns, non-significant; ^*^, p<0.05; ^**^, p<0.01; ^***^, p<0.001; ^****^, p<0.0001 N_cells_ = 66,764. N_spots_ = 170,138. **(D)** Scheme of RecBCD bound by the Gam protein of phage *λ* (subunits S1 and S2). **(E)** RecB spot lifetime histograms (bars) for cells over-expressing the Gam protein, fitted with a bi-exponential decay model (black line, fit components shown as dashed lines). N_cells_ = 8,812. N_spots_ = 18,698. **(F)** Proportion of long-lived RecB spots as a function of ciprofloxacin concentration for cells over-expressing the Gam protein. Violin plots show single-cell distributions, box plots show the median and interquartile range. Statistically significant differences to the WT, no ciprofloxacin condition (Mann-Whitney test) are shown as follows: ns, non-significant; ^*^, p<0.05; ^**^, p<0.01; ^***^, p<0.001; ^****^, p<0.0001 N_cells_ = 8,812. N_spots_ = 18,698.

### RecB forms long-lived complexes with DSBs

Using the time-lapse data, we built histograms of RecB spot lifetimes at the different ciprofloxacin concentrations (Figure 2B). In the absence of ciprofloxacin, the majority of spots lasted less than 2 seconds. Under increasing ciprofloxacin concentrations, the distribution showed a clear shift towards longer-lived spots, with the histogram at 20 and 30 ng/ml ciprofloxacin displaying a clear tail of long-lived events (*>* 10 sec). Fitting these histograms with a mono-exponential decay model (*y* = *a*.*e*^−*k*.*t*^) did not match the experimental data accurately (Supp. Figure 7). In particular, it did not account for the tail of longer-lived spots that formed at high ciprofloxacin concentrations. Fitting with a biexponential decay model 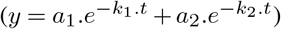 accounted better for the longer-lived RecB spots (Figure 2B). We made sure that the better fit was not solely a result of the increased number of fit parameters by computing Akaike’s Information Criteria (AIC). The average AIC values for each ciprofloxacin concentration (Supp. Table 4) confirmed that the additional parameters helped describe the data better. The bi-exponential fit outlined two populations of spots with different lifetimes: a short-lived one, with average lifetimes ranging from 1.4 to 1.7 sec; and a longer-lived one, with average lifetimes ranging from 10 to 14 sec (Supp. Table 5). Even though the short-lived spots always represented a majority of events (over 90% of the spots), the proportion of long-lived spots increased under higher ciprofloxacin exposure from 1.4 ± 0.3% at 3 ng/ml ciprofloxacin to 5.5 ± 1.4% at 30 ng/ml ciprofloxacin (Figure 2C and Supp. Table 5).

To determine whether short- or long-lived spots resulted from RecB binding to DSBs, we measured RecB spot lifetime in the presence and absence of ciprofloxacin while over-expressing the Gam protein of phage *λ* from a plasmid. The Gam protein was previously shown to bind to the RecBCD complex by mimicking DNA double-strand ends (55) (Figure 2D), and its overexpression is, therefore, expected to prevent RecBCD binding to DSBs. Accordingly, cells that expressed Gam and were exposed to high ciprofloxacin (30 ng/ml) showed little elongation compared to cells that did not overexpress Gam (Supp. Figure 8), indicating that most cells did not induce the SOS response, as a result of the inability of the RecBCD-Gam complex to bind to DSBs and load RecA. The resulting RecB spot lifetimes showed a similar distribution to the one obtained in the absence of ciprofloxacin (Supp. Figure 9). This was confirmed by fitting the histogram with our bi-exponential decay model (Figure 2E), which found a proportion of long-lived spots in the presence of 30 ng/ml ciprofloxacin (1.8 ± 0.9%, Figure 2F and Supp. table 5) equivalent to wild-type cells that were not exposed to ciprofloxacin (1.8 ± 1%). The residual amount of long-lived RecB spots in the presence of Gam could be a result of residual DSB binding despite the presence of Gam. This residual binding could also explain the small increase in cell length observed in Gam-expressing cells under 30 ng/ml ciprofloxacin (Supp. Figure 8). Since Gam overexpression prevents RecB binding to DSBs, and specifically caused long-lived spots to disappear, we conclude that long-lived spots correspond to RecB molecules bound to a DSB (and we will refer to them as DSB-bound RecB throughout this article). Given that under Gam overexpression, 95% of RecB spots had a lifetime shorter than 10 sec (Supp. Figure 10), we considered any spot with a lifetime over 10 sec to be a DSB-bound RecB molecule. The appearance of short-lived spots in the presence of Gam is likely a result of a combination of short-lived DNA interactions and slow, confined diffusion of the RecBCD-Gam-Halo complex (Supp. Note 2 and Supp. Figure 11).

Using the Gam protein allowed us to identify DSB-bound RecB molecules as the longer-lived population in our bi-exponential fits of the RecB spot lifetime histograms. We determined that when it processes a DSB, RecB stays bound to it for 10 to 14 seconds on average, regardless of the ciprofloxacin concentration (Supp. Table 5). This is consistent with processing of DSBs by RecBCD taking place independently of the number of DSBs in the cell. In addition, we utilised the total duration of the experiment (75 minutes) to investigate whether the duration of ciprofloxacin exposure affected spot lifetimes. We performed separate fits at 15 min intervals for cells exposed to 30 ng/ml ciprofloxacin (Supp. Figure 12) and compared spot lifetimes. At all time points, the RecB spot lifetimes obtained were consistent with those reported in Supp. Table 5 (10–15 sec for long-lived spots, and 1.5–2 sec for short-lived spots), indicating that RecB binding was not affected by the duration of ciprofloxacin exposure. This suggests that, as expected, the presence of additional damage due to continuous exposure to ciprofloxacin does not influence the processing of each individual DSB by RecBCD, in keeping with independent processing of each break.

### RecB dissociation time depends on its exonuclease activity

Since the duration of the RecB-DSB interaction was unaffected by the amount of DNA damage, we wondered if it was an intrinsic property of RecBCD processing, or if dissociation of the complex was triggered by the following step in the repair process: the loading of RecA onto DNA. To assess this, we used the Δ*recA* mutant, which lacks the RecA protein, and the *recB*_1080_ mutant, where RecB has lost its exonuclease activity, as well as its RecA-loading activity. We imaged both strains in the presence and absence of ciprofloxacin (30 ng/ml), computed ∼ histograms of RecB spot lifetimes and fitted them with a bi-exponential decay model (Supp. Figure 13), from which we extracted the binding times of RecB on DSBs (Figure 3A and Supp. Table 6). In the Δ*recA* mutant, the DSB-binding lifetime of RecB was similar to the wild-type (~12 sec), both in the presence and absence of ciprofloxacin. This suggests that RecA loading by RecBCD does not play a role in triggering the dissociation of wild-type RecBCD from DNA. In contrast, under exposure to 30 ng/ml ciprofloxacin, RecB_1080_ stayed bound to DSBs for longer on average than wild-type RecB (19.4 ± 7.1 sec), highlighting the importance of its exonuclease activity in controlling its capacity to dissociate from DNA. This could be a result of the DNA strands threading through the RecBCD complex: the wild-type RecBCD digests the DNA strands as it translocates, which likely makes it easier to dissociate, for example, through a short back-tracking motion. In contrast, RecB_1080_ unwinds the DNA strands without digesting them, which would cause significant steric hindrance to dissociation.

**Fig. 3.**
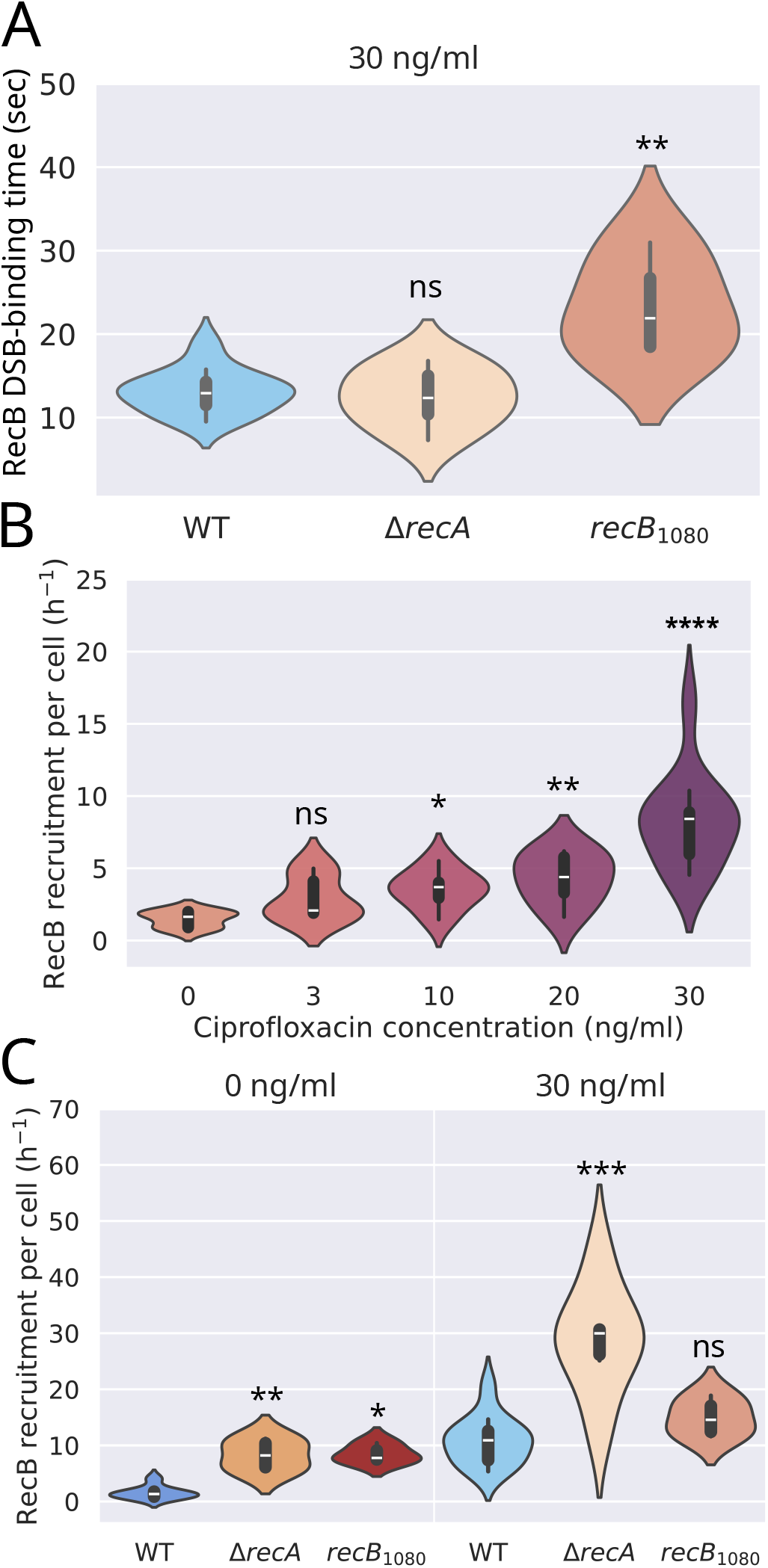
RecB binding to DSBs. **(A)** Lifetime of DSB-bound RecB in wild-type (WT) cells, and in the Δ*recA* and *recB*_1080_ mutants. Statistically significant differences are shown relative to wild-type cells exposed to 30 ng/ml ciprofloxacin. N_cells_ = 32,427. N_spots_ = 116,763. **(B)** Recruitment rate of RecB to DSBs per cell at different ciprofloxacin concentrations. Statistically significant differences are shown relative to the no ciprofloxacin condition. N_cells_ = 66,764. N_spots_ = 170,138. **(C)** Recruitment rate of RecB to DSBs per cell for the Δ*recA* and *recB*_1080_ mutants. N_cells_ = 55,843. N_spots_ = 175,600. For all panels, violin plots show single-cell distributions, box plots show the median and interquartile range. p-values from a Mann-Whitney test are shown as follows: ns, non-significant; ^*^, p<0.05; ^**^, p<0.01; ^***^, p<0.001; ^****^, p<0.0001

### RecB binding can be used to estimate the rate of DSB formation

Whereas the lifetime of RecB spots informs us on how long RecB stays bound to DSBs, the rate of appearance of spots can be used to estimate the rate of recruitment of RecB to DSBs under each DNA damage condition. To calculate this rate, we used the RecB spot lifetime histogram fits to estimate the total number of slow-dissociating spots per unit of time. Since we determined that the appearance of a slow-dissociating spot corresponds to the binding of RecB to a DSB, we could calculate the number of RecB recruitments to DSBs per cell per hour (Figure 3B). We estimated that 1.3 ±1.1 RecB molecules are recruited to DSBs per hour in the absence of ciprofloxacin. As expected, the recruitment rate of RecB increased with ciprofloxacin concentration, up to 8.2 ±3.6 RecB recruited to DSBs per hour at 30 ng/ml ciprofloxacin.

In the Δ*recA* and *recB*_1080_ mutants, the rate of initial DSB formation is expected to be the same as in the wild-type, as these mutations only affect the repair process once DSBs are present. Surprisingly, the rate of recruitment of RecB to DSBs in the Δ*recA* mutant was higher than in the wild-type, both in the absence (1.6 ± 1.1 *h*^−1^ and 8.3 ± 2.8 *h*^−1^) and the presence (10.8 ± 4.3 *h*^−1^ and 28.8 ± 9.5 *h*^−1^) of ciprofloxacin (Figure 3C). We hypothesise that this higher rate of RecB recruitment to DSBs is a result of multiple recruitment events on the same original DSB that cannot be repaired because of the absence of RecA. In the *recB*_1080_ mutant, RecB recruitment was higher than in the wild-type in the absence of ciprofloxacin (1.6 ± 1.1 *h*^−1^ and 8.5 ± 1.7 *h*^−1^), and equivalent to WT cells in the presence of ciprofloxacin (Figure 3C). This might reflect the ability of the *recB*_1080_ mutant to repair DSBs albeit with lower efficiency than wild-type cells (see Discussion).

### Ciprofloxacin exposure induces formation of RecA filaments

During DSB processing, RecBCD facilitates the loading of the RecA protein onto ssDNA. To broaden our view of the repair process downstream of RecBCD, we imaged a tandem fusion of RecA with the fluorescent protein SYFP2, which was previously shown to preserve RecA functionality (9). Based on the fluorescence distribution in the cells (Supp. Figure 14A), we identified three states of RecA: diffuse; forming a bright focus; or forming an elongated structure (called “filament” thereafter), which matches previous *in vivo* observations of RecA (9). Because of the large diversity of shapes observed, particularly for RecA filaments, detecting RecA structures using rule-based algorithms was challenging. Therefore, we designed and trained a deep-learning algorithm capable of classifying individual cells based on the three types of spatial distributions cited above (Supp. Figures 4 and 14B). In cells not exposed to ciprofloxacin, the RecA-associated fluorescence was mostly diffuse, and in ∼20% of the cells it formed either a bright focus or a filament. This is consistent with the expectation that RecA diffuses freely in the cell without DNA damage, and polymerises on DNA in the event of an endogenous DSB. After one hour of exposure, the proportion of cells that contained a RecA filament had increased to ∼30% at 20 ng/ml ciprofloxacin, and ∼45% at 30 ng/ml. RecA foci, on the other hand, formed rapidly following ciprofloxacin exposure (present in ∼40% of the cells after 15 min of exposure to 30 ng/ml ciprofloxacin) and dissipated after ∼1 hour. Taken together, these results show that RecA foci are transient structures in the repair process that do not accumulate under high DNA damage, whereas RecA filaments accumulate under constant exposure to ciprofloxacin. These results are consistent with previous imaging of RecA (9, 42, 44, 45), which have reported transient foci when DSBs occur at a replication fork and can be repaired rapidly (42), but longer-lived filaments upon exogenous DNA damage (9, 44, 45).

### DSB-bound RecB colocalises with the bacterial nucleoid

RecA loading onto ssDNA triggers the SOS response, which inhibits cell division, leading to cell filamentation. Previous studies have reported that the SOS response also triggers compaction of the bacterial nucleoid (23). To see if this was the case for breaks induced by ciprofloxacin in our experimental conditions, we stained DNA using the Sytox Green dye. The use of a green dye allowed us to image RecB concomitantly, and to correlate the position of DSB-bound RecB molecules (defined as RecB spots with a lifetime *>*10 sec, Supp. Figure 10) with that of the nucleoid. In cells that were not exposed to ciprofloxacin, the nucleoid was often observed to be bi-lobed (Figure 4A), consistent with our previous observations (38). After 60 min of exposure to ciprofloxacin, the cells appeared elongated, and the nucleoid was often compacted in the centre of the cell. As a result of nucleoid compaction, the total fraction of the cell occupied by the nucleoid decreased, from 40% on average in untreated cells to 31% in cells exposed to 30 ng/ml of ciprofloxacin for one hour (Supp. Figure 15). In addition to the compaction, we observed that nucleoids were increasingly centred, and the cells elongated upon increasing exposure to ciprofloxacin, both in time and concentration (Supp. Figure 16).

**Fig. 4.**
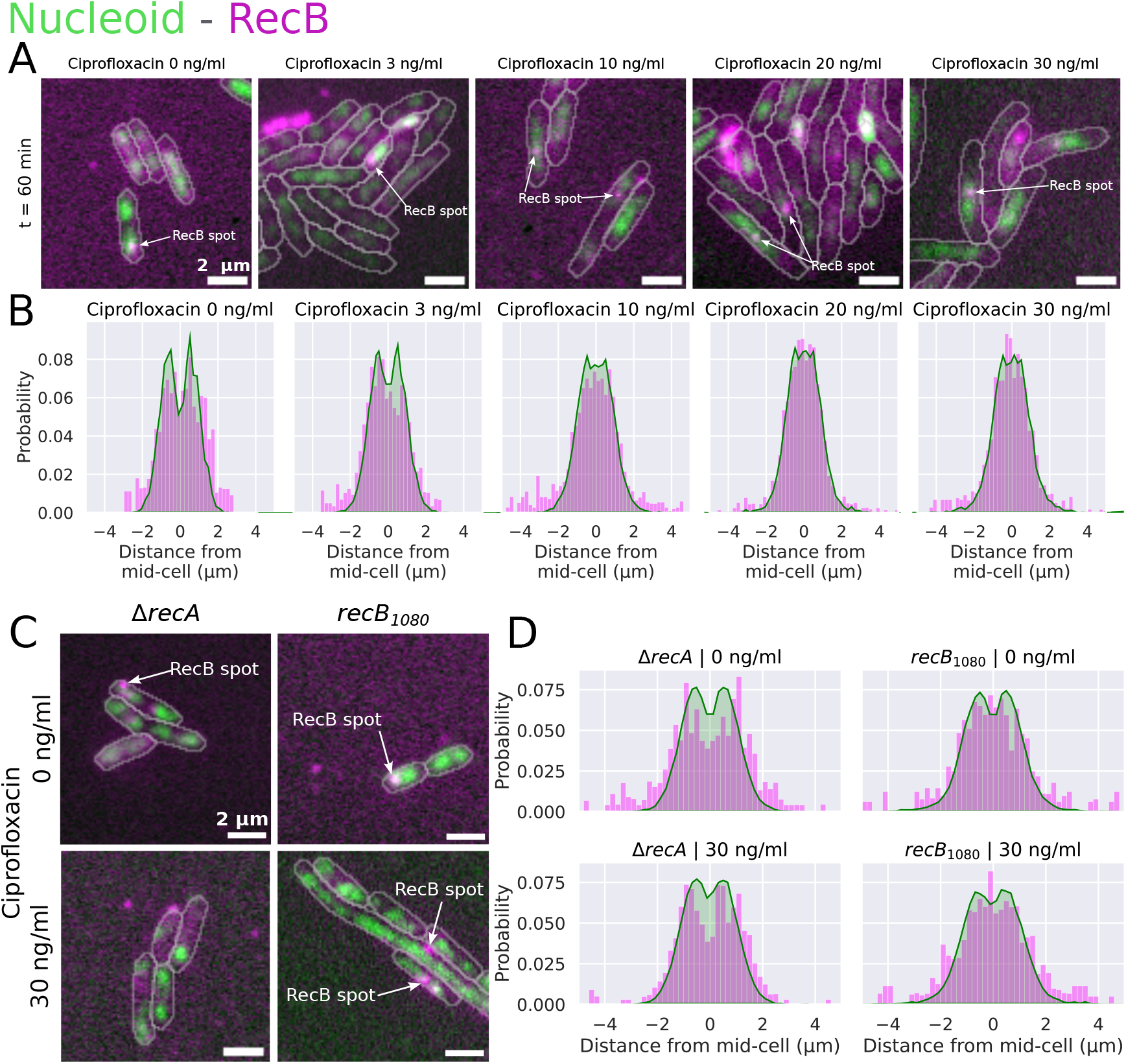
Colocalisation of RecB spots with the bacterial nucleoid. **(A)** Representative images of cells (segmented outline in grey) showing the nucleoid (green) and RecB-associated fluorescence (magenta). RecB spots (indicated by arrows) are located in close proximity to the nucleoid. **(B)** Overlay of nucleoid density (green area) and position of DSB-bound RecB molecules (magenta bars) along the cell’s long axis, for different ciprofloxacin concentrations (0 to 30 ng/ml). N_cells_ = 24,014. N_spots_ = 79,969. N_nucleoids_ = 31,441. **(C)** Representative overlay images of RecB-associated fluorescence (magenta) and the bacterial nucleoid (green) in the Δ*recA* and *recB*_1080_ mutants. Segmented cell outline shown in grey. **(D)** Overlay of nucleoid density (green area) and position of DSB-bound RecB molecules (magenta bars) along the cell’s long axis for the Δ*recA* and *recB*_1080_ mutants at 0 and 30 ng/ml ciprofloxacin. N_cells_ = 7,804. N_spots_ = 36,595. N_nucleoids_ = 11,215.

Long-lived RecB spots were found close to the nucleoid (Figures 4A and 4B), thus strengthening our interpretation that long-lived spots are DSB-bound RecB molecules. This proximity was observed at any time during the experiment (Supp. Figure 17), suggesting that it does not depend on the number of DSBs present in the cell. Upon exposure to increasing ciprofloxacin concentrations, induction of the SOS response caused the cells to elongate significantly (from 3.1 ± 0.19 µm on average in the absence of ciprofloxacin to 5.5 ± 0.71 µm after 75 min of exposure to 30 ng/ml ciprofloxacin, Supp. Figure 8). Despite this, and because of compaction and centring, the nucleoid remained within ∼2 µm either side of the cell centre. The distribution of DSB-bound RecB molecules overlapped strongly with nucleoid density (Figure 4B), indicating that as the nucleoid was compacted, additional DSB formation triggered RecB recruitment at the centre of the cell, where the nucleoid was located.

### Nucleoid compaction requires RecA loading

The spatial distribution of nucleoid density and DSB-bound RecB was affected in the Δ*recA* and *recB*_1080_ mutants (Figures 4C and 4D). In the Δ*recA* mutant, exposure to ciprofloxacin did not trigger nucleoid compaction and centring as in wild-type cells. This confirmed that nucleoid compaction under ciprofloxacin exposure requires RecA loading, and most likely the induction of the SOS response. Despite the absence of compaction, the nucleoid occupied a smaller fraction of the cell in the Δ*recA* mutant after exposure to ciprofloxacin (Supp. Figure 18). This can be attributed to the progressive degradation of the bacterial chromosome by the repeated cycles of RecBCD binding (Figure 3C). In the *recB*_1080_ mutant, the nucleoid often formed irregular shapes and small segregated regions in the cell (Figure 4C). This disorganisation was especially visible in the presence of 30 ng/ml ciprofloxacin. Furthermore, in the *recB*_1080_ mutant the nucleoid did not undergo compaction within the 75 min of our experiment (Supp. Figure 18). This might reflect the less efficient RecA loading in this mutant, leading to different time dynamics of the SOS induction and either impaired or significantly delayed nucleoid compaction.

In both mutant strains, the localisation of DSB-bound RecB overlapped with the bacterial nucleoid (Figure 4D), similar to the wild-type. In the Δ*recA* mutant, addition of 30 ng/ml ciprofloxacin did not change the spatial distribution of nucleoid density or DSB-bound RecB. This result was expected, as Δ*recA* cells are unable to induce the SOS response, and, therefore, do not undergo nucleoid compaction. In the *recB*_1080_ mutant in the absence of ciprofloxacin, both nucleoid density and DSB-bound RecB distributions were more centred than in wild-type cells, despite the nucleoid occupying the same fraction of the cell area (Supp. Figure 18). This might be because of SOS induction in this mutant, which occurs even in the absence of DNA damaging agent (38). Upon exposure of the *recB*_1080_ mutant to ciprofloxacin, the absence of change in the distribution of nucleoid density and DSB-bound RecB position reinforced the hypothesis that nucleoid compaction is modified in this mutant.

## Discussion

### RecB dissociation does not depend on subsequent repair steps

In this work, we used single-molecule imaging of RecB to elucidate key aspects of RecBCD’s processing of DSBs *in vivo*. Deep-learning-based quantitative image analysis allowed us to analyse a very large number of cells (in total 98,828, with at least 380 per experiment) and to infer how long RecB remains bound to DNA after DSB recognition, and whether this binding time depends on the level of DNA damage. We found that in wild-type *E. coli*, RecB stays bound to DSBs for 10 to 15 seconds on average (Supp. Table 5), independent of the concentration of ciprofloxacin. Given the high speed of RecBCD exonuclease activity *in vivo* [1.6kb/s (3)], this would translate into an end-resection process that encompasses on average 20–30 kb of the chromosome, in keeping with what has been observed previously (3, 5). Additionally, we investigated whether RecA might play a role in promoting RecBCD dissociation from DNA. Imaging RecB in a Δ*recA* mutant indicated that this is not the case, and that RecB dissociation occurs independently of RecA loading (Table 6). Interestingly, the *recB*_1080_ mutant had a reduced dissociation rate compared to wild-type cells, which highlights the importance of RecB’s exonuclease activity in the process and the possibility that digestion of the unwound DNA strands before Chi recognition is a pre-requisite for RecBCD dissociation.

### RecB can be used as a proxy to quantify DSB formation *in vivo*

Given RecBCD’s high affinity for DSBs *in vivo* and our ability to detect its binding to DNA ends, the system is a useful tool to detect the presence of DSBs in *E. coli*. Our experiments allowed us to estimate the number of recruitments of RecB to DSBs per hour, under different ciprofloxacin concentrations (Figure 3B), and in the Δ*recA* and *recB*_1080_ mutants (Figure 3C). In the absence of ciprofloxacin, we estimated that 1.3 ±1.1 RecB molecules are recruited to DSBs per hour. A previous study had estimated that 18% of cells generated an endogenous DNA double-strand end per cell cycle (12), for a cell cycle length of 20 min (shorter than the 35 min cell cycle time in our experiments). This would correspond to a rate of ∼0.6 DSBs per hour, a similar value but slightly lower than our estimate. This discrepancy could be due to the significant differences in the methods and growth conditions used to produce these estimates.

In the wild-type strain, the general pathway for DSB repair suggests that RecBCD is recruited once per DSB (Figure 5). We can, therefore, reasonably expect that the observed rate of RecB recruitments matches the number of DSBs formed (1 to 8 per hour, depending on ciprofloxacin concentration). One should, however, note that because RecBCD abundance is low (47), this system can only be used at a relatively low concentration of antibiotic, as a very high number of DSBs would saturate the

**Fig. 5.**
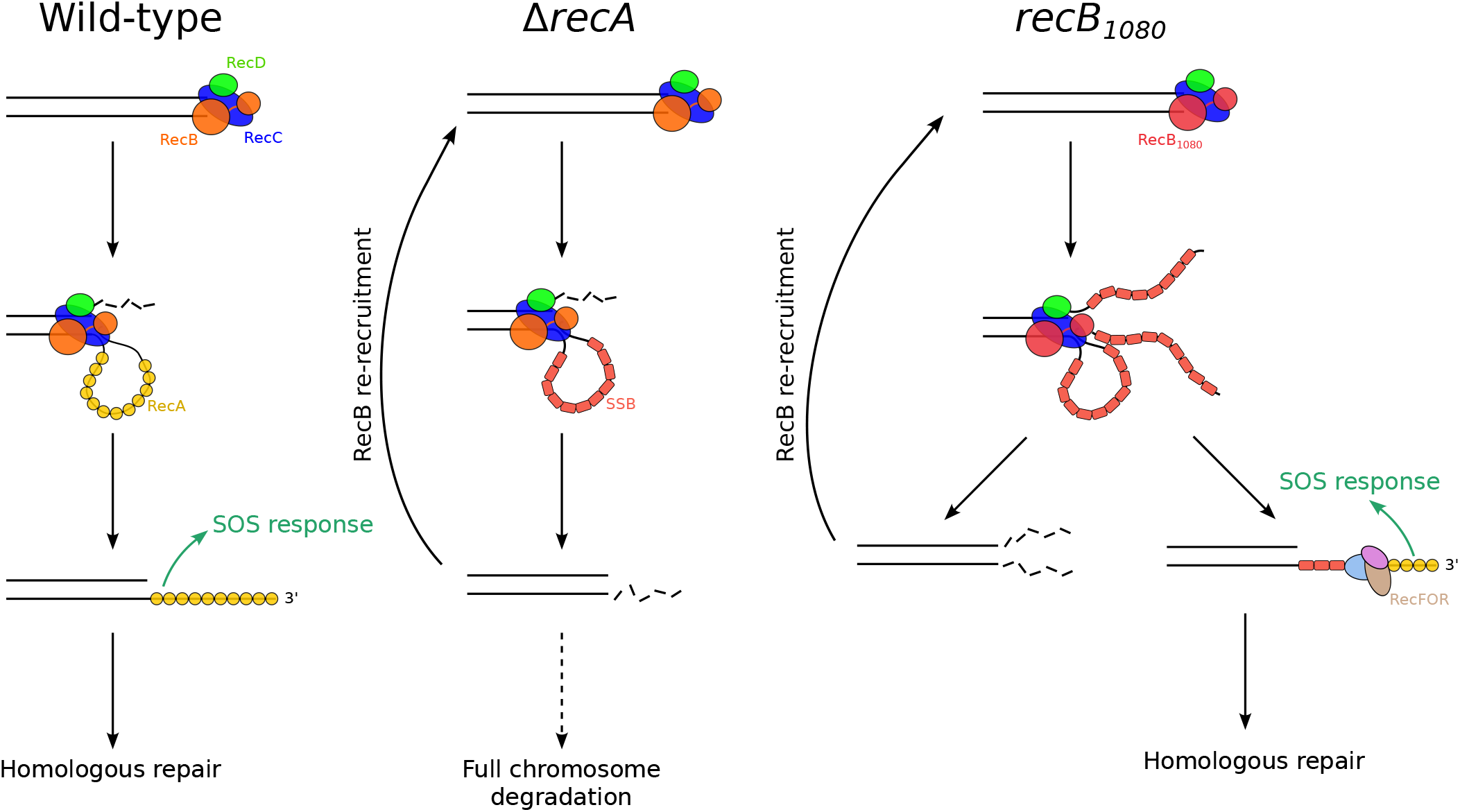
RecBCD recruitment pathways in wild-type *E. coli*, Δ*recA* and *recB*_1080_ mutants. **(Wild-type)** After DSB recognition, RecBCD degrades DNA until it recognises a Chi-site. It switches activity to create a 3’ ssDNA overhang, and promotes RecA loading. The RecA-coated ssDNA can then be used for DNA repair by homologous recombination. **(Δreca)** In the absence of RecA, the 3’ ssDNA is coated by SSB, and eventually digested by cellular nucleases. Blunting of the DNA end by digestion of the ssDNA creates a new substrate for binding of RecBCD. **(RecB**_**1080**_ ) Following DSB recognition, RecB_1080_ unwinds DNA without digesting it. The unwound ssDNA can either be digested by nucleases, leading to a new blunt dsDNA end and RecBCD re-recruitment; or the RecFOR complex displaces SSB to load RecA, allowing DNA repair by homologous recombination to proceed.

RecBCD complex. In the Δ*recA* mutant, the DSB repair pathway disruptions lead to multiple RecB recruitments per DSB and, therefore, this system cannot be used to estimate DSB formation rates reliably.

It should be noted that in some cases, ciprofloxacin is expected to produce double-sided DSBs, hence, leading to two RecB recruitments per break. We did not take this into account in our estimation of the DSB formation rate, as it has been previously suggested that the two sides of a DSB are kept in close proximity during DNA repair (56, 57), and our imaging setup would not allow us to separate two RecB spots located close together. Indeed, our timelapse images of RecB binding seldom show two spots next to each other, further supporting the idea that we cannot resolve RecBCD binding to both sides of a DSB.

Imaging DSBs in live cells is challenging and was previously achieved using the Gam protein of bacteriophage Mu, which binds to free double-stranded DNA ends (58–60). Indeed, treatment with 30ng/ml ciprofloxacin was reported to create ∼5 Gam foci per cell in keeping with the ∼8 DSB/cell/hour we observe. However, Gam binding to DSBs prevents RecBCD action and, therefore, the repair of the breaks. Using RecBCD as a probe allows us to monitor DSB formation in live *E. coli* in real-time, with single-cell sensitivity and without perturbing the repair process. However, one limitation of using the Halo-tag for labelling is that it does not allow for long-term imaging (over several hours), as the labelled protein progressively becomes replaced with newly synthesised, unlabelled protein. This tool could, therefore, be further improved by substituting the Halo tag with a bright and photostable fluorescent protein [as was used in our previous work (47)], enabling long-term imaging, for example, in a mother-machine microfluidic device.

### RecB is recruited to DSBs multiple times in the Δ*recA* mutant

Our observations of RecB recruitment to DSBs (Figures 3B and 3C) and nucleoid position (Figures 4C and 4D) in the different mutant strains have led us to the model of RecB recruitment described in Figure 5, which confirms the model we previously proposed (38), and extends it by proposing that RecBCD is reloaded multiple times on the same DSB. In wild-type cells, a DSB is recognised by RecBCD, which promotes RecA loading. The RecA filament triggers the SOS response, and is used for homology search and repair. In the Δ*recA* mutant, the 3’ ssDNA generated by RecBCD is first coated by SSB, and then degraded by the SbcCD and ExoI nucleases (61). This leads to blunting of the DNA end, creating a new substrate on which RecBCD can bind. This cycle likely leads to multiple RecBCD recruitments per DSB, and eventually to full chromosome degradation. In the *recB*_1080_ mutant, RecBCD unwinds DNA without degrading it. Our data show that at equal levels of DNA damage induction, RecB_1080_ is recruited to the DSBs less frequently than RecBCD in Δ*recA* cells (Figure 3C, 8.5 ± 1.7 for RecB_1080_ and 28.8 ± 9.5 for Δ*recA*). This suggests that in the *recB*_1080_ mutant, two competing pathways take place: either DNA degradation by SbcCD and ExoI leading to DNA-end blunting and re-recruitment of RecBCD; or RecA loading by RecFOR, leading to SOS induction and homologous repair. Thus, the RecFOR-dependent pathway would provide an escape route from the cycle of re-recruitment.

### RecA filaments accumulate under high DNA damage

Under exposure to high ciprofloxacin concentrations (20–30 ng/ml), we observed an accumulation of cells that contained a RecA filament (Supp. Figures 14A and 14B). At these ciprofloxacin concentrations, we have determined that cells undergo frequent DSBs, leading to multiple recruitments of RecB to DSBs over the course of the experiment (up to 8 per hour, Figure 3B). Given RecBCD’s high processivity (3), such a high number of RecB recruitments would likely result in several tens of kilobases of DNA being digested at different chromosomal locations, which could result in the complete absence of a homologous copy of the DSB site. In this case, we expect that the repair process will stall at the homology search stage after formation of the RecA filament, which is consistent with our observation that a large proportion of cells contain RecA filaments following exposure to high ciprofloxacin concentrations.

### RecB colocalises with the bacterial nucleoid

Imaging RecB simulteanously to the bacterial nucleoid reinforced our interpretation that long-lived RecB spots (*>*10 sec) are DSB-bound RecB molecules, and helped us understand their spatial distribution in the cell. As expected, DSB-bound RecB molecules colocalised with the nucleoid. Nucleoid compaction due to SOS induction and RecN activity had previously been reported after UV irradiation and treatment with mitomycin C (23, 56), and we were able to confirm that a similar compaction occurs under ciprofloxacin treatment (Supp. Figure 15). In the Δ*recA* mutant, the lack of nucleoid compaction suggests the involvement of RecA loading and SOS induction in nucleoid compaction, again confirming previous observations (23, 56). In the *recB*_1080_ mutant, the nucleoid is more centred than in the wild-type strain but we did not observe any additional compaction upon exposure to ciprofloxacin. This could result from a substantial delay in SOS induction and recN expression as a result of the loading of RecA by the RecFOR alternative pathway (38).

Taken together, this work provides a detailed picture of RecBCD’s interaction with DNA following DSBs. We found that RecB dissociation is independent of RecA but depends on RecB’s exonuclease activity, and that binding time is unaffected by DNA damage levels. Additionally, we observed changes in nucleoid conformation in response to DNA damage and examined the effects of high antibiotic levels on the formation of RecA filaments. The model we developed offers an overview of RecBCD recruitment to DSBs and dissociation from DNA, depending on the strain’s ability to effectively carry out or complete repair. These findings contribute valuable insights into the dynamics of DNA repair mechanisms in *E. coli*, suggesting that RecBCD dissociation from DNA is an intrinsic property of the complex, largely unaffected by external factors, such as DNA damage levels or RecA loading.

## ACKNOWLEDGEMENTS

We would like to thank Prof. David Leach for insightful discussions on DNA repair in *E. coli*. We thank Prof. Mark Dillingham for sharing his knowledge on the biochemistry of RecBCD, and for gifting the Gam plasmid. We thank Prof. Johan Elf for kindly sharing the RecA-SYFP2 tandem fusion strain. We thank Dr. Jean Ollion for the development of BACMMAN and its associated tools, as well as the image analysis support he provided. We thank Elise Darmon for discussing results, and reviewing the manuscript.

## AUTHOR CONTRIBUTIONS

M.E.K. and D.T. conceived the experiments and designed the data analysis. D.T., A.L., and L.McL. built the strains. D.T. collected and analysed the data. M.E.K., D.T. and A.L. discussed the data. D.T. wrote the manuscript’s first draft and created the figures. M.E.K., D.T. and A.L. revised the manuscript. L.McL., as the lab manager, oversaw order placements and maintained the lab organisation. M.E.K. provided funding. All authors read, edited and approved the final manuscript.

