## Supplementary information for "Spatio-temporal coordination of the DNA Double-Strand Break repair machinery in *Escherichia coli*"

### Single molecule imaging of RecB *in vivo* reveals dynamics of DNA Double Strand Break repair in *Escherichia coli*

#### Supplementary Note 1: JF549 dye photobleaching kinetics

In our approach to estimating RecB binding times to DSBs, the photostability of the fluorescent dye was of primary concern. Since we detect single molecules of RecB bound to DSBs, two main photophysical behaviours were liable to cause artefacts in our data:

1. Photobleaching (an irreversible loss of fluorescence) would limit the maximum number of frames we can track a DSB-bound RecB for, and would artefactually reduce the fitted binding times since some fluorescent spots might disappear as a result of photobleaching rather than unbinding from DNA

2. Blinking (a transient loss of fluorescence) could cause repeated disappearance and reappearance of single molecules, and, hence, bias the computed binding times and recruitment rates to DNA.

To address these concerns, we computed ensemble-level photobleaching curves by integrating the total fluorescence signal from the cells (Supp. Fig. 2A). After 50 frames of laser exposure, a significant amount of photobleaching was noticeable (27% $\pm$ 9 of the initial fluorescence remaining). The exact photobleaching rate of the dye in our experimental conditions can be estimated by fitting the photobleaching curve with an exponential decay function of the form $y = a.e^{-k.t} + b$ (with a the amplitude of the fit, k the bleaching rate, and b an offset to account for cellular auto-fluorescence). Since the bleaching rate is expected to mostly depend on experimental parameters that vary between but not within datasets (output laser power, HiLo angle), one bleaching rate was computed per dataset (Supp. Fig. 2B). The true RecB dissociation rates were computed by subtracting the corresponding bleaching rates from the fitted spot disappearance rates.
All experimental photobleaching curves were well-fitted by a mono-exponential decay function. If significant blinking of the dye was occurring, we would expect photobleaching curves to follow more complex kinetics (62). We therefore concluded that in our experimental conditions, the JF549 dye was not experiencing blinking, consistent with previous reports of the dye's outstanding photostability (63).

#### Supplementary Note 2: Formation of RecB spots by the freely diffusing RecBCD-Halo-Gam complex

In cells that overexpress the Gam protein, RecB is expected to be unable to bind DSBs. Even though Gam overexpression prevents the apparition of long-lived RecB spots under exposure to ciprofloxacin, short-lived RecB spots remain present (Figure 2E, Supp. Figure 9 and Supp. Table 5). To understand how RecB spots can be formed in the absence of DSB binding, we simulated the expected displacements of a RecBCD-Halo-Gam complex according to the distribution of single displacements for a random 2D walk:

$$P(r) = \frac{r}{2D_i t} e^{-\frac{r^2}{4D_i t}} \quad \textbf{(2)}$$

with $D_i$ the apparent diffusion coefficient, and $t$ the frame time. Our previous work found a diffusion coefficient for RecB-Halo of $\sim$1.5 µm$^2$.s$^{-1}$ (38). Assuming that the diffusion coefficient of the complex (that does not interact with DSBs) is proportional to the cube of the molecular weight, we expect a diffusion coefficient for the full RecBCD-Halo-Gam complex (386 kDa, 353 kDa without the Gam protein) of $\sim$1 µm$^2$.s$^{-1}$. The frame time in our experiments is 1 second, but it is likely that a molecule that would stay immobile for a fraction of that time would still be visible as a spot. Although calculating a precise value is challenging, we used a ballpark estimate of 500 ms (half our frame time) of a molecule being immobile to be detected as a spot in our experiment. Supp. Figure 11 shows the distribution of expected displacements under these parameters. Although most displacements would be too large to result in a bright fluorescent spot, a small fraction ($\sim$4%) are smaller than 300 nm, and could create a spot. When factoring in the number of RecB molecules per cell [5 on average (47)] and the number of frames in our timelapse (50), this would result in $\sim$10 RecB spots per cell, on the same order of magnitude as the number of spots observed in our experiments in the presence of the Gam protein (4.1 $\pm$ 0.2, mean $\pm$ sem). Note that a single RecB molecule can form several spots over the length of the timelapse experiment.

**Supplementary Table 1.** List of bacterial strains used in this study

| Name | Description | Reference |
|---|---|---|
| MEK2623 | MG1655 *recB::halotag recA::syfp2* | this work |
| MEK2622 | MG1655 *recB::halotag* Δ*recA* | this work |
| MEK716 | MG1655 *recB1080::halotag* | this work |
| MEK2324 | MG1655 *recB1080::halotag HK022::psfiA-GFP* | (38) |
| MEK65 | MG1655 *recB::halotag* | (47) |
| MEK2629 | MG1655 *recB::halotag pBad::GamL* | this work |
| DL654 | MG1655 Δ*recA* | (64) |

**Supplementary Table 2.** List of bacterial plasmids used in this study

| Name | Description | Reference |
|---|---|---|
| pSF1 | Expression of HaloTag under the control of the pBAD promoter | (47) |
| pDT6 | GamL inserted into pBAD322K by restriction cloning | (55) |

**Supplementary Table 3.** Acquisition parameters used for microscopy.

| Channel | Illumination | Intensity | Exposure (ms) | EM gain | Images (interval) | Z-stack |
|---|---|---|---|---|---|---|
| Brightfield | Lamp | NA | 30 | 0 | 1 | 16 slices, 0.2 µm step |
| JF549 | 561-nm | 2 mW | 1000 | 150 | 50 (2 sec) | No |
| SYFP2 | 488-nm | 2 mW | 50 | 100 | 50 (2sec) | No |
| Sytox Green | 488-nm | 2 mW | 50 | 100 | 1 | No |

**Supplementary Table 4.** Akaike's Information Criteria (AIC) for mono- and bi-exponential decay fits of the RecB spot lifetime histograms. Lower values (in bold) show the most relevant model. Values are given as the mean ± standard deviation on all datasets. Diff. shows the difference in mean AIC value between the two models (negative values are in favour of the bi-exponential model).

| Ciprofloxacin concentration (ng/ml) | AIC (mono-exponential fit) | AIC (bi-exponential fit) | Diff. |
|---|---|---|---|
| 0 ng/ml | -258.5 ± 117.0 | **-284.5** ± 140.1 | -26 |
| 3 ng/ml | -307.0 ± 36.6 | **-322.2** ± 42.3 | -15 |
| 10 ng/ml | -290.4 ± 48.3 | **-340.2** ± 75.3 | -50 |
| 20 ng/ml | -286.4 ± 45.0 | **-349.6** ± 49.1 | -63 |
| 30 ng/ml | -303.5 ± 40.3 | **-387.5** ± 62.0 | -84 |

**Supplementary Table 5.** Parameters derived from the spot lifetime histogram fits (Figures 2B and 2C). The lifetime was calculated as the inverse of the fitted dissociation rate. Values are given as the median ± standard deviation over at least 3 independent datasets. $N_{cells}$ = 66,764. $N_{spots}$ = 170,138

| Strain | Cipro. | Type | Lifetime (sec) | Population (%) |
|---|---|---|---|---|
| WT | 0 ng/ml | Short | 1.4 ± 0.2 | 98.2 ± 1.0 |
| | | Long | 9.5 ± 9.7 | 1.8 ± 1.0 |
| | 3 ng/ml | Short | 1.5 ± 0.1 | 98.6 ± 0.3 |
| | | Long | 10.5 ± 1.3 | 1.4 ± 0.3 |
| | 10 ng/ml | Short | 1.7 ± 0.2 | 97.6 ± 0.6 |
| | | Long | 13.7 ± 2.3 | 2.4 ± 0.6 |
| | 20 ng/ml | Short | 1.6 ± 0.2 | 96.6 ± 0.8 |
| | | Long | 11.8 ± 2.1 | 3.4 ± 0.8 |
| | 30 ng/ml | Short | 1.7 ± 0.2 | 94.5 ± 1.4 |
| | | Long | 13.3 ± 2.2 | 5.5 ± 1.4 |
| Gam | 0 ng/ml | Short | 1.7 ± 0.2 | 99.2 ± 0.4 |
| | | Long | 15.5 ± 9.1 | 0.8 ± 0.4 |
| | 30 ng/ml | Short | 1.5 ± 0.1 | 98.2 ± 0.9 |
| | | Long | 8.2 ± 1.8 | 1.8 ± 0.9 |

**Supplementary Table 6.** Parameters derived from the spot lifetime histogram fits for the Δ*recA* and $recB_{1080}$ strains (Supp. Figure 13). The lifetime was calculated as the inverse of the fitted dissociation rate. Values are given as the median ± standard deviation over at least 3 independent datasets. $N_{cells}$ = 56,131. $N_{spots}$ = 177,646.

| Strain | Cipro. | Type | Lifetime (sec) | Population (%) |
|---|---|---|---|---|
| Δ*recA* | 0 ng/ml | Short | 1.8 ± 0.2 | 96.6 ± 1.0 |
| | | Long | 13.0 ± 1.8 | 3.5 ± 1.0 |
| | 30 ng/ml | Short | 2.1 ± 0.2 | 90.5 ± 3.4 |
| | | Long | 12.4 ± 3.5 | 9.5 ± 3.4 |
| $recB_{1080}$ | 0 ng/ml | Short | 1.8 ± 0.2 | 97.4 ± 0.9 |
| | | Long | 16.7 ± 5.5 | 2.6 ± 0.9 |
| | 30 ng/ml | Short | 2.1 ± 0.1 | 96.5 ± 0.7 |
| | | Long | 19.4 ± 7.1 | 3.5 ± 0.7 |

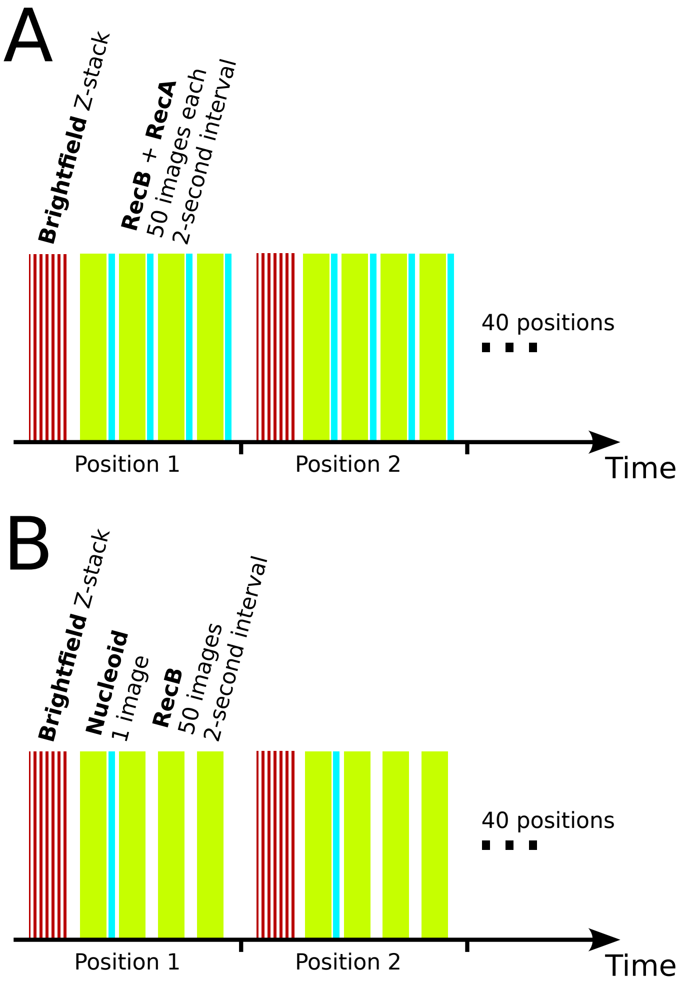


**Supplementary Figure 1.** Acquisition patterns for microscopy experiments. **(A)** Acquisition pattern for experiments with brightfield, RecB (JF549) and RecA (SYFP2) channels. **(B)** Acquisition pattern for experiments with brightfield, RecB (JF549) and Nucleoid (Sytox Green) channels.

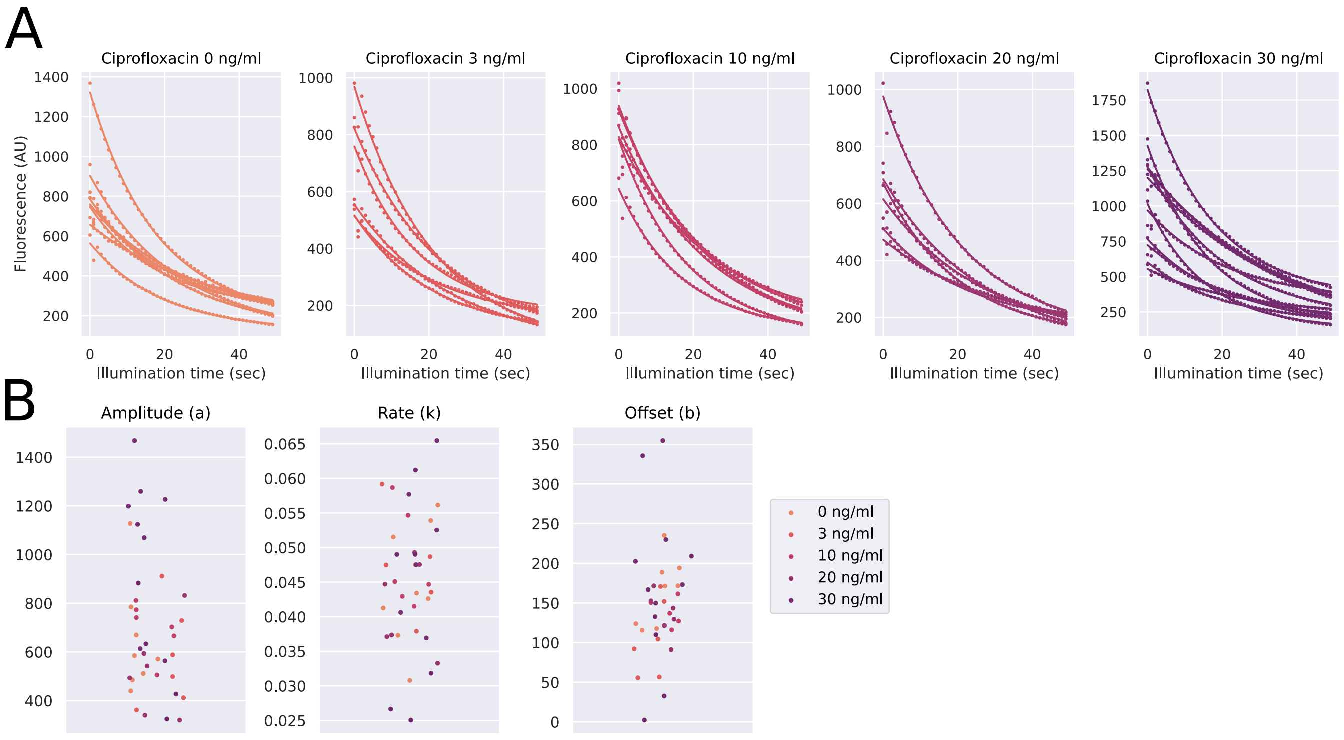


**Supplementary Figure 2.** Ensemble-level photobleaching of the JF549 dye. **(A)** Average background-subtracted fluorescence for independent datasets (dots), overlaid with the photobleaching rate fit ($y = a.e^{-k.t} + b$, line). $N_{cells}$ = 66,764. **(B)** Fitted model parameters for each dataset: amplitude (a), photobleaching rate (k) and offset (b). $N_{cells}$ = 66,764.

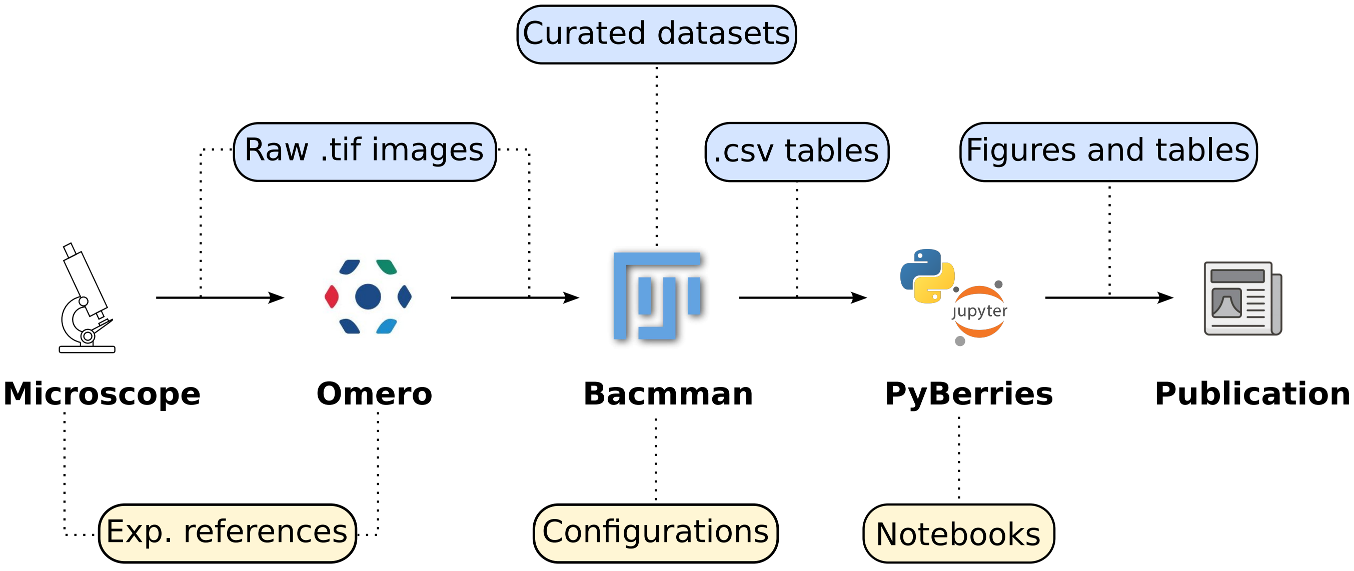


**Supplementary Figure 3.** Data storage and analysis pipeline used in this study. Blue labels indicate stored data and yellow labels indicate code and references that would allow reproducing the different analysis steps.

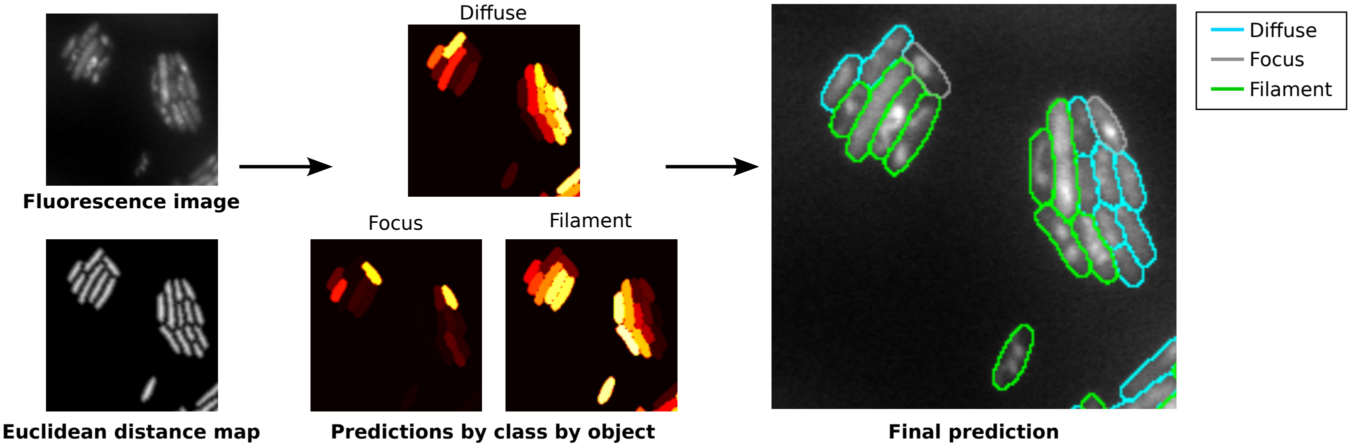


**Supplementary Figure 4.** Classification of cells according to the RecA structures they contain by our in-house Unet-based deep-learning network.

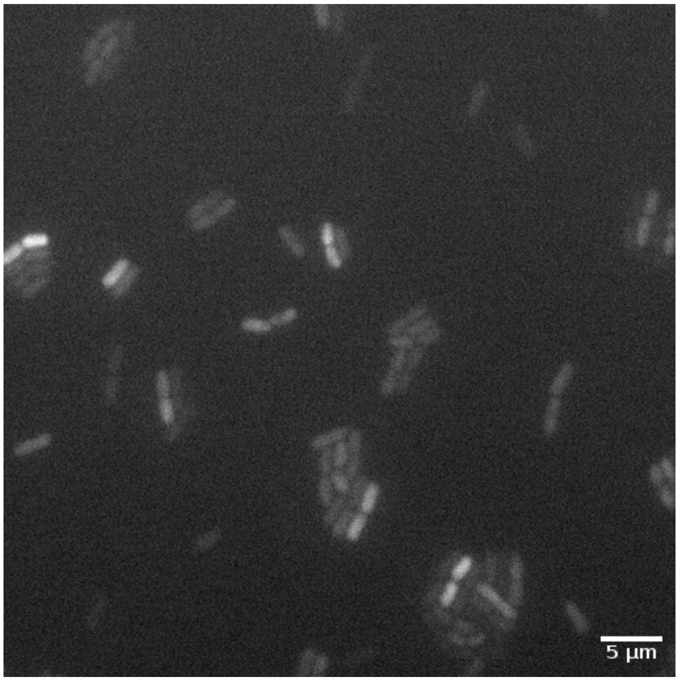


**Supplementary Figure 5.** Representative fluorescence image (1 second exposure time) of freely diffusing Halo-tag expressed from a pBAD plasmid (induced with 1% w/v arabinose) in MG1655 *E. coli* cells.

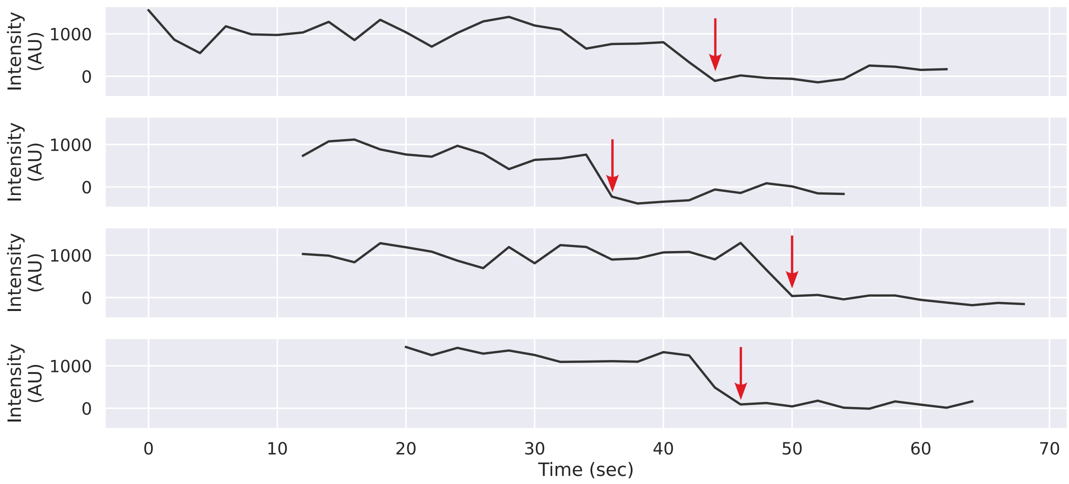


**Supplementary Figure 6.** Representative background-subtracted intensity time traces (black lines) for single RecB spots, showing loss of intensity in a single step (red arrows).

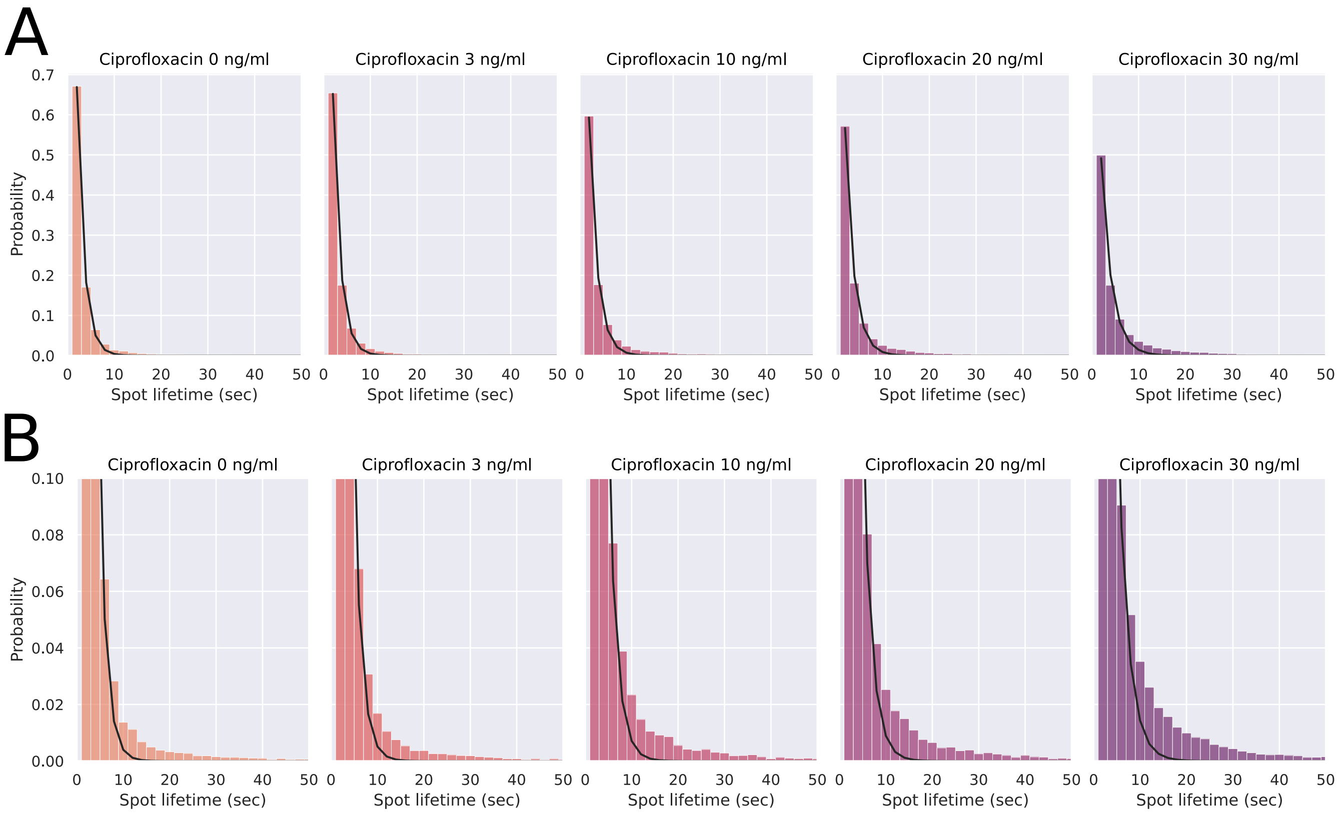


**Supplementary Figure 7. (A)** Histograms of RecB spot lifetime (bars) under exposure to ciprofloxacin, with overlaid mono-exponential decay fits ($y = a.e^{-k.t}$, black line). $N_{cells}$ = 66,764. $N_{spots}$ = 170,138 **(B)** Zoom on the tails of histograms in (A).

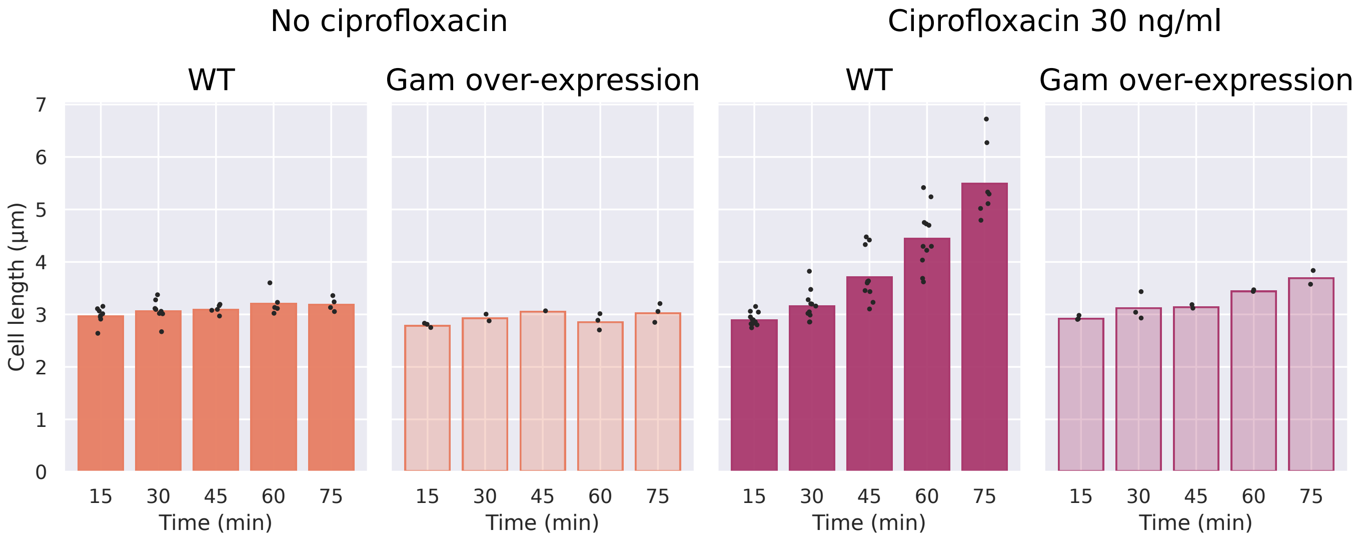


**Supplementary Figure 8.** Average length of cells that over-express Gam or not (WT), under exposure to 0 or 30 ng/mL ciprofloxacin. Black dots show averages for indvidual datasets, and bars the average between them. $N_{cells}$ = 41,403.

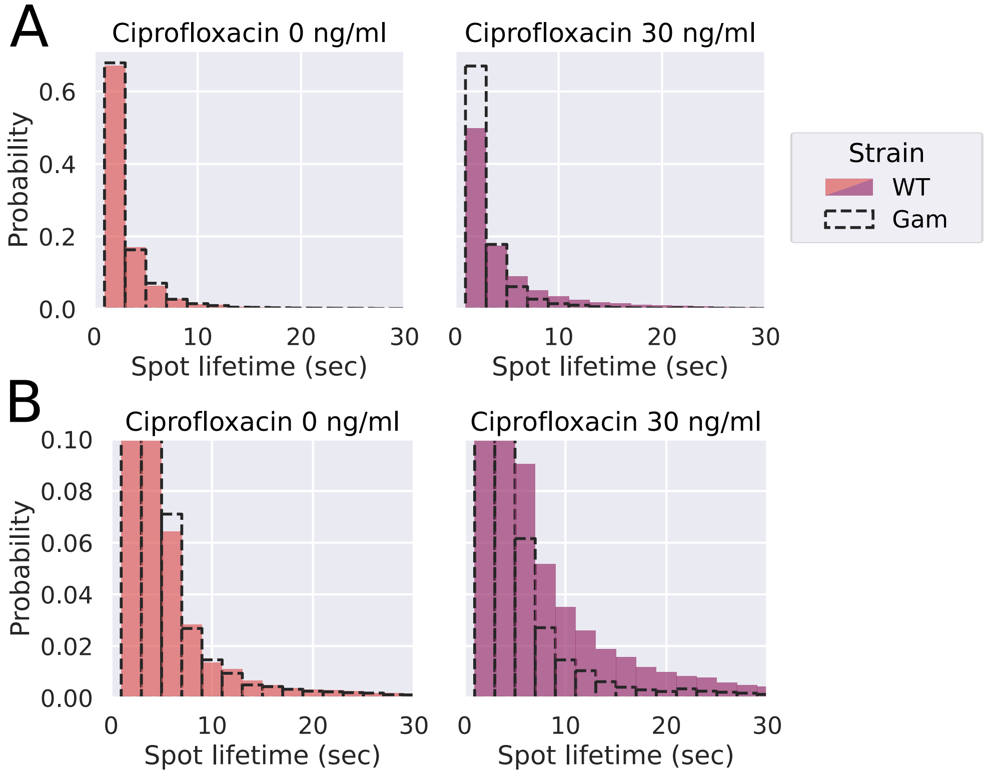


**Supplementary Figure 9. (A)** Histograms of RecB spot lifetimes for wild-type cells (coloured bars) and cells overexpressing Gam (dashes). $N_{cells}$ = 41,403. $N_{spots}$ = 96,453 **(B)** Zoom on the tail of the histograms from (A).

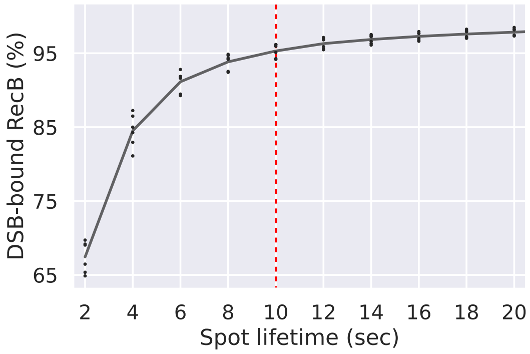


**Supplementary Figure 10.** Proportion of DSB-bound RecB molecules according to RecB spot lifetime. Black dots show averages for individual datasets; the black line is the average between them, and the red dashed line shows the smallest lifetime at which RecB spots have a 95% probability of being DSB-bound. $N_{cells}$ = 8,812. $N_{spots}$ = 18,698.

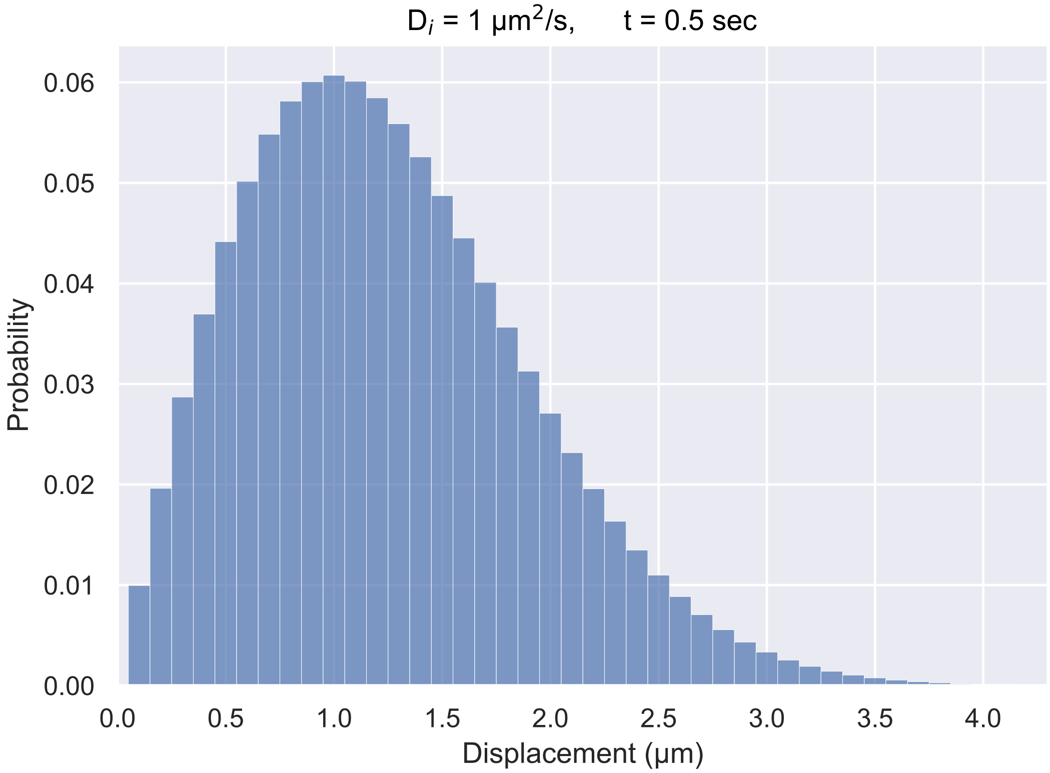


**Supplementary Figure 11.** Histogram of expected displacements for a molecule diffusing at 1 $\mu m^2.s^{-1}$ over a 500 ms frame time.

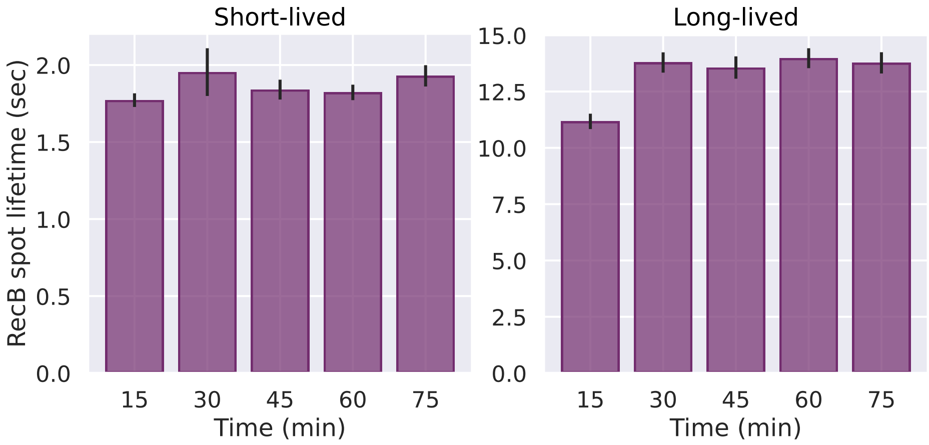


**Supplementary Figure 12.** Fitted lifetimes of short- and long-lived RecB spots, following different durations of exposure to 30 ng/ml ciprofloxacin. Coloured bars represent the fitted lifetimes, and black strokes the standard error of the mean obtained by bootstrapping. $N_{cells}$ = 18,945. $N_{spots}$ = 54,475

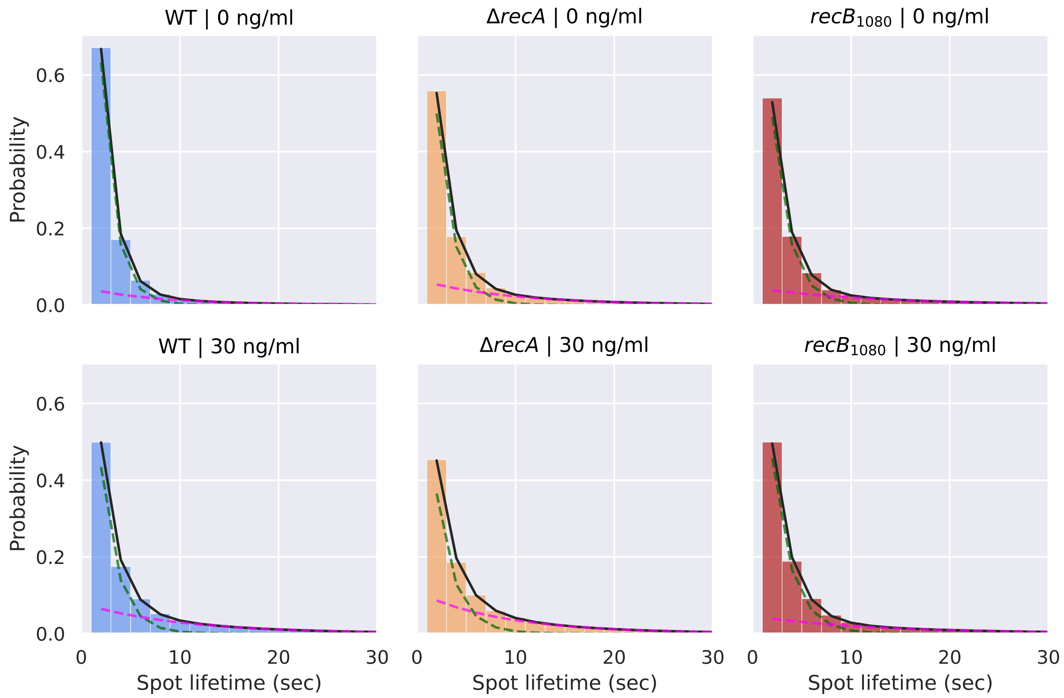


**Supplementary Figure 13.** RecB spot lifetime histograms for wild-type (WT), $\Delta$*recA* and *recB*$_{1080}$ mutants, at 0 and 30 ng/mL ciprofloxacin, fitted with a bi-exponential decay model (black line, fit components showed as dashed lines). $N_{cells}$ = 56,131. $N_{spots}$ = 177,646.

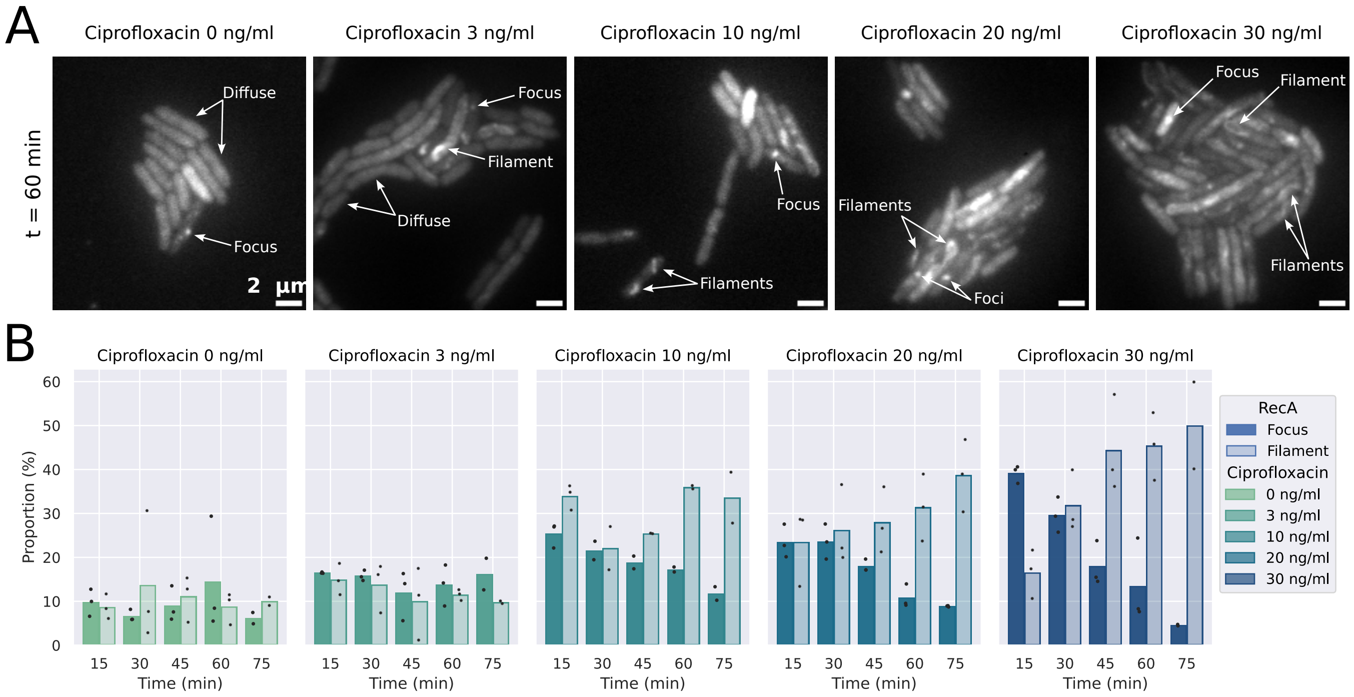


**Supplementary Figure 14.** RecA structures formed upon exposure to ciprofloxacin. **(A)** Representative images of cells containing different RecA structures (diffuse fluorescence, foci or filaments) after 60 minutes of exposure to ciprofloxacin. Arrows point to representative examples of each of these structures. **(B)** Proportion of cells containing RecA foci or filaments. Black dots represent averages for individual datasets, and bars the average between them. $N_{cells}$ = 32,031.

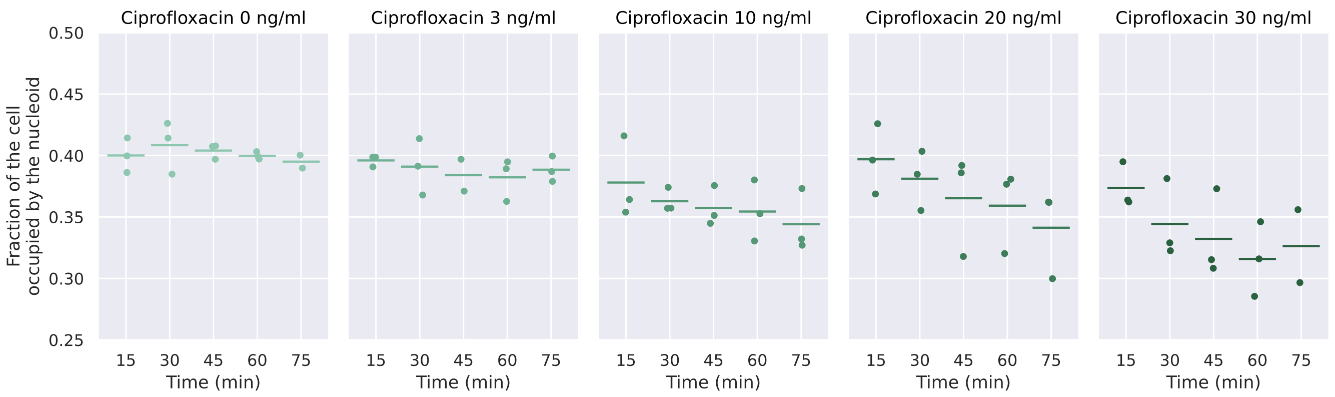


**Supplementary Figure 15.** Average fraction of the bacterial cell occupied by the nucleoid (stained using the Sytox Green dye) at different ciprofloxacin concentrations (0 to 30 ng/ml) and duration of exposure (15 to 75 min). Dots represent averages for individual datasets, and dashes the average between them. $N_{cells}$ = 24,014. $N_{nucleoids}$ = 31,441.

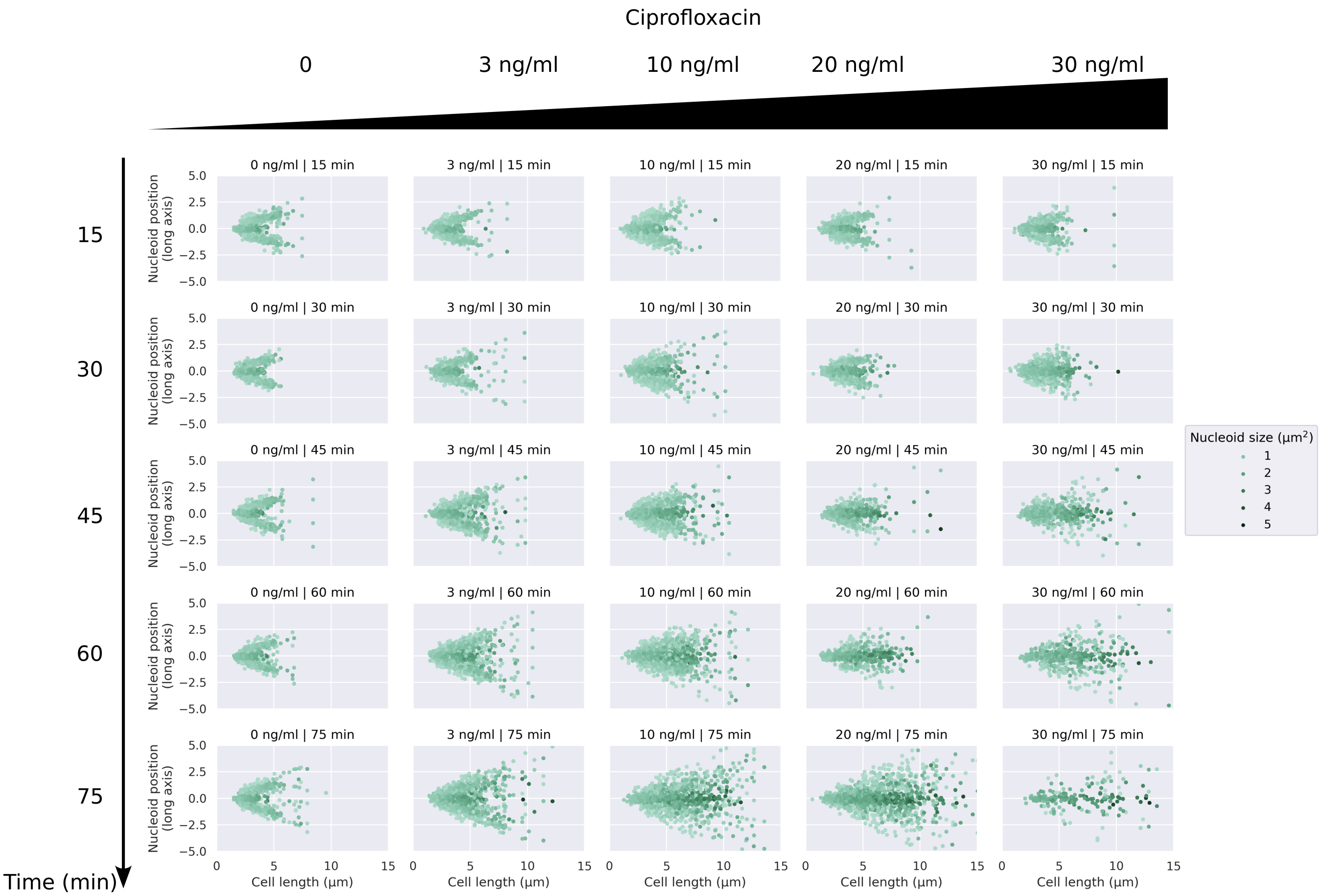


**Supplementary Figure 16.** Position of the nucleoid along the cell's long axis against cell length for different ciprofloxacin concentrations (columns) and durations of ciprofloxacin exposure (rows). Point colour indicates the total surface covered by the nucleoid in the cell, in $\mu m^2$. $N_{cells}$ = 24,014. $N_{nucleoids}$ = 31,441.

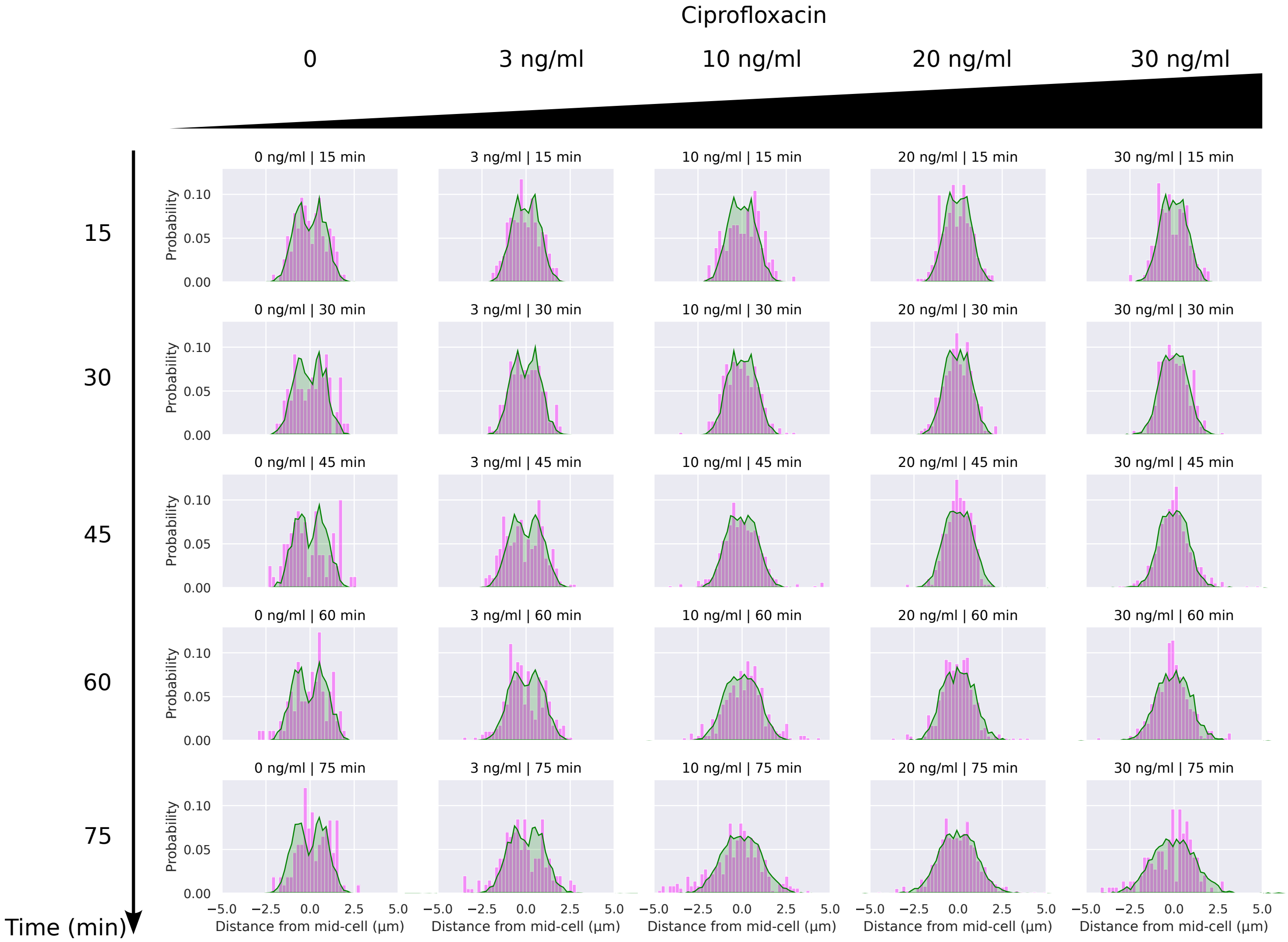


**Supplementary Figure 17.** Overlay of nucleoid density (green area) and position of DSB-bound RecB molecules (magenta bars) along the cell's long axis, for different ciprofloxacin concentrations (columns) and durations of exposure (rows). $N_{cells}$ = 15,029. $N_{spots}$ = 58,331. $N_{nucleoids}$ = 20,831.

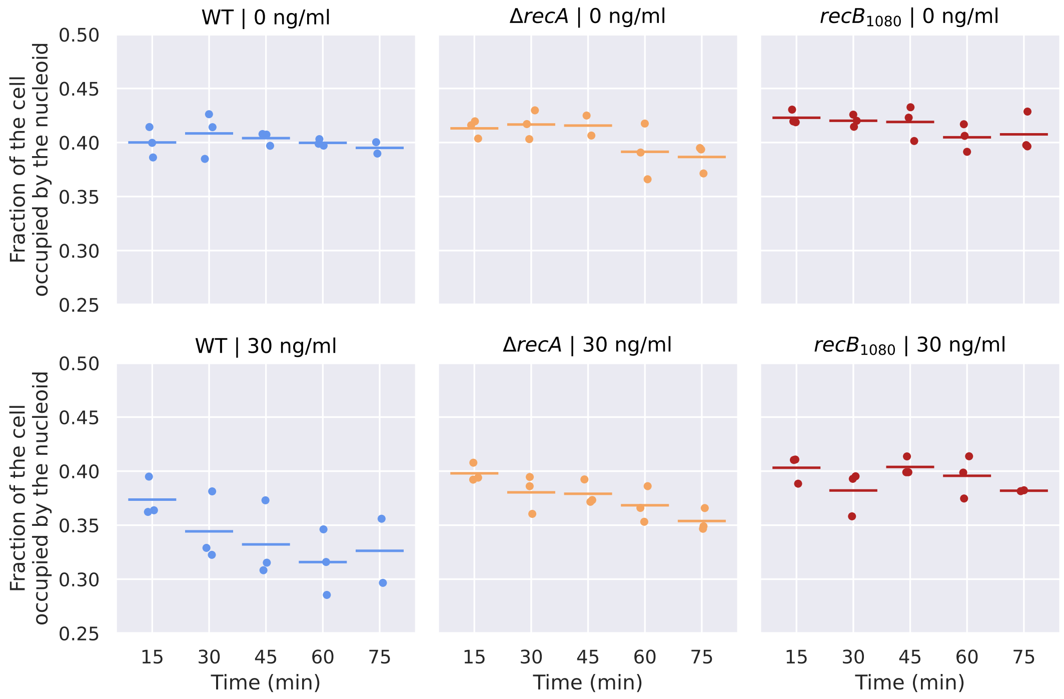


**Supplementary Figure 18.** Average fraction of the bacterial cell occupied by the nucleoid (stained using the Sytox Green dye) at different ciprofloxacin concentrations (0 to 30 ng/mL) and duration of exposure (15 to 75 min), for wild-type cells (reproduced from Supp. Figure 15 for comparison) and the $\Delta$*recA* and *recB*$_{1080}$ mutants. Dots represent averages for individual datasets, and dashes the average between them. $N_{cells}$ = 15,029. $N_{nucleoids}$ = 20,831.
